# Ductal myofibroblasts reactivate contractile program to stabilize alveolar architecture during lung regeneration

**DOI:** 10.64898/2026.09.23.753969

**Authors:** Hiroaki Katsura, Akira Yamaoka, Naozane Nomura, Junhyeong Kim, Osamu Nishimura, Dooseon Cho, Mitsutaka Kadota, Takefumi Kondo, Takaya Abe, Hiroshi Kiyonari, Daisuke Hazama, Shinya Tane, Yoshimasa Maniwa, Tatsuya Nagano, Mitsuru Morimoto

## Abstract

The alveolar sac architecture is essential for efficient gas exchange and must be precisely maintained throughout life; however, how this delicate structure is preserved during adult regeneration remains poorly understood. Using a mouse pneumonectomy model, we found that *Lgr6*+ *Hhip*+ ductal myofibroblasts, a poorly characterized mesenchymal population, are indispensable for maintaining alveolar integrity during lung regrowth. Comprehensive characterization using single-cell transcriptomics, mouse genetics, and pharmacological assays demonstrated that these ductal myofibroblasts secrete myogenic factors, most notably CCN4, to reactivate a myogenic program that converts them into contractile PA-DMFs, thereby preserving alveolar architecture. Lineage-tracing further revealed that these ductal myofibroblasts originate from embryonic MCAM- SMA+ distal progenitors via subepithelial TGF-β signaling, serving as a lifelong guardian of alveolar structural integrity. Notably, cross-species analysis identified an analogous population of *LGR6*+ fibromyocytes in human respiratory bronchioles. Together, these findings indicate ductal myofibroblasts as a developmentally programmed cell population that reactivate a contractile program to structurally support the regeneration of adult lungs.

## Introduction

Lung tissue architecture is fundamentally coupled with respiratory function. Consequently, disruption of this architecture often leads to pulmonary diseases, such as bronchopulmonary dysplasia (BPD) and chronic obstructive pulmonary disease (COPD). Under these conditions, irreversible structural damage to the alveoli leads to impaired gas exchange and progressive decline in respiratory function. BPD, a condition that primarily affects premature infants, is characterized by arrested alveolar development, leading to simplified alveolar structures^1^. In contrast, COPD involves progressive destruction of the alveolar walls in adults, leading to enlarged airspaces and reduced elastic recoil^2^. Although the developmental processes of alveolar septation and maturation have been extensively studied, the mechanisms underlying alveolar structural regeneration in the adult lungs remain poorly understood. Addressing this gap is essential for elucidating the pathogenesis of chronic lung diseases and developing therapeutics to restore alveolar integrity.

The alveolar tissue architecture is established during postnatal alveologenesis, a process in which immature sac-like structures are subdivided into a highly organized network of mature alveoli to enable efficient gas exchange^3,4^. During this period, two types of smooth muscle actin-positive (SMA+) mesenchymal populations are present in the distal lungs: alveolar myofibroblasts and ductal myofibroblasts. Although both cell types express SMA, they originate from distinct molecular lineages and exhibit different spatial and temporal dynamics during postnatal lung development. Alveolar myofibroblasts transiently emerge in the distal lung and drive alveolar septation via contractile forces associated with myosin light chain phosphorylation^5–11^. These cells are abundant in alveoli during the first 2 weeks after birth, begin to disappear shortly after the completion of alveologenesis, and are largely absent from adult lungs. Their differentiation is regulated by signaling pathways including platelet-derived growth factor A (PDGF-A)^12^, Sonic hedgehog (SHH)^13^, and transforming growth factor-β (TGF-β)^14^. Consequently, the disruption of these pathways impairs alveolarization and results in simplified alveolar structures, which are hallmarks of developmental lung diseases. In contrast, ductal myofibroblasts are specifically localized in alveolar ducts, which are transitional structures that connect the airways and alveoli. Unlike alveolar myofibroblasts, ductal myofibroblasts are maintained beyond postnatal development and reside in adult lungs. Moreover, recent single-cell RNA sequencing (scRNA-seq) studies have identified ductal myofibroblasts as a distinct mesenchymal population in the developing lung^15–17^; however, their physiological roles in alveologenesis and adult lung regeneration remain poorly defined. Simultaneously, extracellular matrix (ECM) components, particularly elastin fibers, provide essential mechanical support for alveolar integrity^18^. Hence, impaired elastogenesis leads to severe structural abnormalities^19–23^. Although these studies have determined the individual importance of myofibroblasts and the ECM in alveologenesis, the spatiotemporal coordination between these components that shapes the alveolar architecture remains poorly understood.

Therefore, in this study, we employed a compensatory lung growth model following pneumonectomy (PNX) to investigate alveolar structural regeneration in adult lungs. This approach provides a robust platform for studying lung regeneration in the absence of fibrosis or overt tissue injury. In this process, surgical removal of one lung lobe induces rapid expansion of the remaining lobes, leading to the restoration of lung volume and function^24–26^. This regenerative response is characterized by the coordinated proliferation and differentiation of alveolar epithelial cells, particularly type II alveolar epithelial (AT2) cells, which give rise to type I cells that rebuild the gas exchange surface^27,28^. Concomitantly, the alveolar structure undergoes dynamic remodeling, including changes in alveolar size and number, to accommodate the increased mechanical demand^29,30^. Despite these well-described responses, the specific mesenchymal cell populations that contribute to alveolar integrity by supporting and stabilizing regenerating lung tissue remain to be identified.

Accordingly, we identified and characterized a previously underappreciated mesenchymal population located in the alveolar ducts that preserves the alveolar architecture during lung regeneration. Using lineage tracing, spatial transcriptomics, and scRNA-seq, we showed that *Lgr6*⁺ *Hhip*⁺ ductal myofibroblasts represent a resident mesenchymal population established during embryonic development. During compensatory lung regrowth after PNX, ductal myofibroblasts reactivate a contractile programs to stabilize alveolar structures via autocrine myogenic induction. Furthermore, we demonstrated that an analogous population of *LGR6*⁺ fibromyocytes exists in human lungs, suggesting that these structural supporting cells may provide critical insights into the pathology of human pulmonary diseases.

## Results

### Transient emergence of SMA⁺ cells during both development and regeneration in alveoli

Despite their conceptual similarities, previous studies have evaluated developmental and regenerative alveologenesis separately. Therefore, we initially reexamined the progression of alveolarization from postnatal day (P) 1 to one month, providing a detailed spatiotemporal characterization of the developing alveoli. As previously reported, SMA+ alveolar myofibroblasts appear to form a secondary septum structure at the neonatal stage, followed by rapid disappearance beyond two weeks of age (Fig. 1A-B, Supplementary Fig.1A)^6,7,9^. Notably, elastin fiber deposition occurred in spatiotemporal coordination with SMA+ alveolar myofibroblasts, forming a structural scaffold before these cells disappeared. (Fig. 1C-D)^8^. Based on these results, we generated elastogenesis-deficient mice by knocking out the *Fbln5* gene, which is required for proper alveolar architecture maturation (Supplementary Fig.1B–E)^21,22^. Our results revealed an emphysema phenotype in the mutant lungs from P14, whereas no phenotype was observed by P12 (Fig. 1E, Supplementary Fig.1F-G). These observations indicate a structural transition within the first 2–3 weeks of life, shifting from myofibroblast-driven secondary septation during the neonatal period to a more permanent elastin fiber-based scaffold.

**Fig. 1.**
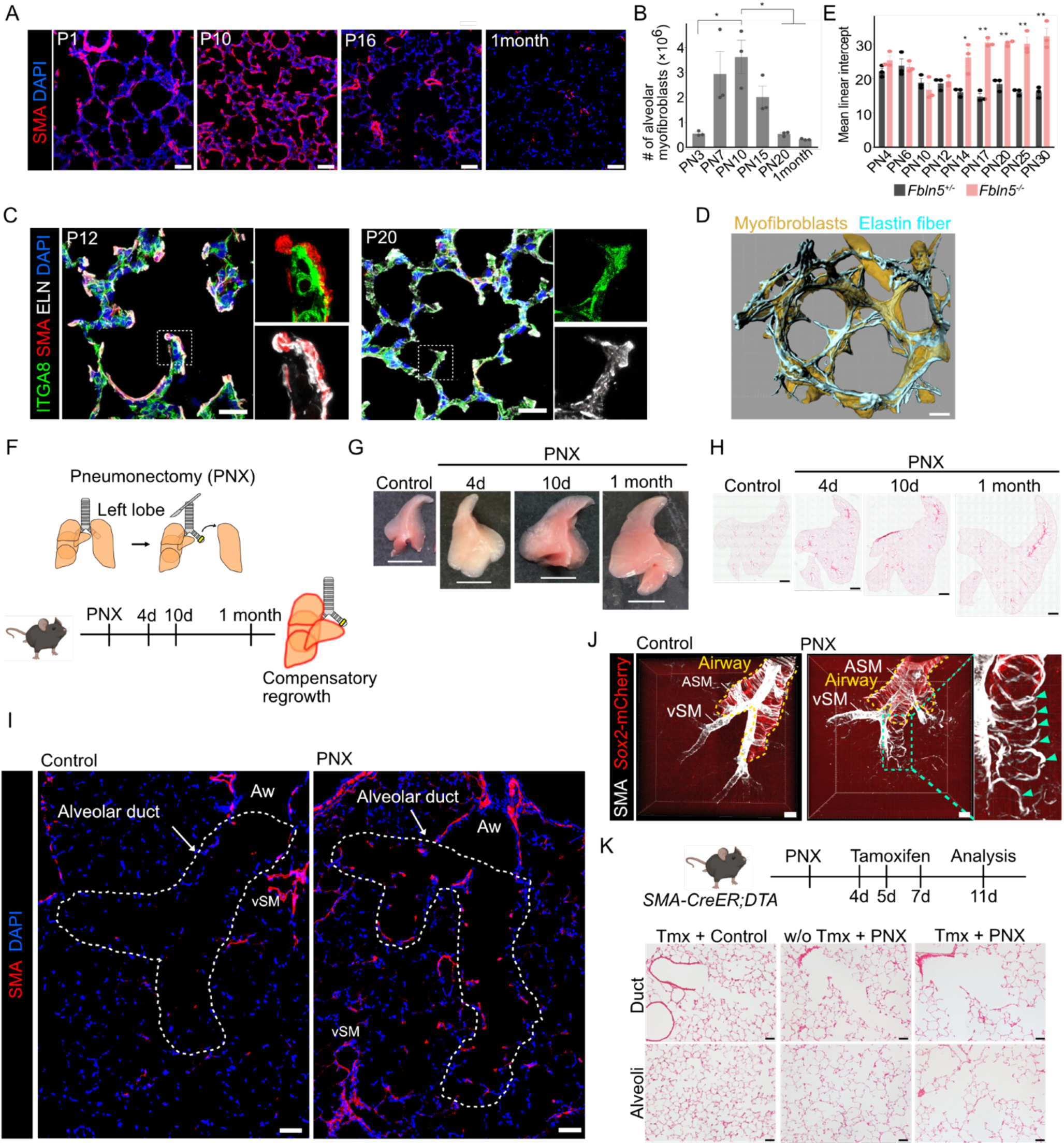
SMA+ cells emerged in the alveoli are required to establish the alveolar structural integrity during development and regeneration. A. Immunofluorescence images of postnatal lung sections stained for SMA. Scale bars, 50 µm. B. Quantification of the number of alveolar myofibroblasts. (n = 3 mice/group) C. Immunofluorescence images of lung sections stained for ITGA8 (green), SMA (red), and ELN (white) at P12 (top) and P20 (bottom). Scale bars, 20µm. D. Three-dimensional reconstruction of myofibroblasts and elastin fibers in alveoli at P10. scale bars, 10 µm. E. Quantification of MLI of *Fbln5*+/- (gray) and *Fbln5*-/- (red) at each time point. (n = 3 mice/group) F. Schematic diagram of time course observation of compensatory lung regrowth after PNX. G. Pictures of accessory lobes at each time point after PNX. Scale bars, 5 mm. H. Time course images of H&E staining of accessory lobes after PNX. Scale bars, 1 mm. I. Immunofluorescence images of lung sections stained for SMA (red) in control (left) and 7d after PNX (right). Scale bars, 50 µm. White dashed lines indicate alveolar duct. Aw, airways. J. Three-dimensional images of cleared lung tissues from *Sox2*-H2B-mCherry reporter mice stained for SMA (white). Yellow dashed line indicates airways and arrowheads indicate the SMA+ cells in the duct. Scale bars, 100 µm. K. Schematic diagram of experimental design to ablate SMA+ cells followed by PNX and images of H&E staining of alveolar ducts (top) and alveoli (bottom) in each condition. Scale bars, 50 µm. All data represent the mean ± SEM obtained from at least three independent mice. \**P*<0.05, \*\**P*<0.01.

We then used a PNX model that triggered compensatory lung growth in the remaining lobes to determine whether similar structural dynamics occur during adult lung regeneration (Fig. 1F-H). Immunofluorescence analysis showed that in control lungs, SMA+ cells were rarely detected in the alveoli, except for in the airway and vascular smooth muscle cells (Fig. 1I, left). However, after PNX, SMA+ cells appeared in the alveolar ducts connecting the terminal bronchioles to the alveoli (Fig. 1I, right). Additionally, three-dimensional analysis revealed that these post-PNX-SMA+ cells were localized as an extension of the airway and formed ring-like structures resembling airway smooth muscles (Fig. 1J, Supplementary Fig.1H). SMA expression was also observed in a subset of pericytes characterized by protrusions attached to blood vessels in the control and PNX lungs (Supplementary Fig.1I-K)^31^. Based on these results, we administered PNX to *SMA-CreER;DTA* mice, in whom SMA-expressing cells were selectively ablated by tamoxifen injection, to determine the functional role of post-PNX-SMA+ cells during regrowth. SMA depletion alone did not cause any detectable structural abnormalities under homeostatic conditions. However, regional emphysematous enlargement of the alveolar ducts and alveoli was observed after PNX (Fig. 1K). These findings indicate that post-PNX-SMA+ cells emerge in the alveolar ducts during lung regrowth and are required to preserve alveolar architecture. Collectively, these results suggest that the transient emergence of SMA+ cells contributes predominantly to the establishment and maintenance of alveolar integrity during development and regeneration.

### *Lgr6*+ ductal myofibroblasts reactivate SMA expression and become post-PNX-SMA+ cells during lung regeneration

We performed lineage-tracing analyses to define the cellular origin of post-PNX-SMA+ cells within lung fibroblast populations. These lung fibroblasts exhibit marked heterogeneity, encompassing multiple distinct cellular subtypes^32–34^. Therefore, we first examined whether adult alveolar fibroblasts contribute to the post-PNX-SMA+ cell population using *Pdgfra-CreER;Ai9* mice, which are widely used to analyze pan-fibroblast populations in the lungs. Immunofluorescence analysis revealed that the *Pdgfra*-lineage cells rarely contributed to the post-PNX-SMA+ cell population (Supplementary Fig. 2A-B) indicating that resident adult alveolar fibroblasts are not the primary source of regenerative myofibroblasts.

Although alveolar myofibroblasts largely disappear after postnatal alveologenesis, recent studies have suggested that a subset of these cells can be maintained until adulthood^7,35^. Thus, we labeled alveolar fibroblasts and myofibroblasts at P1 in *Pdgfra-CreER;Ai9* mice and performed PNX in adulthood to determine whether post-PNX-SMA+ cells originated from postnatal alveolar myofibroblasts. Numerous TOMATO+ alveolar fibroblasts were present in the alveoli; however, most of the post-PNX-SMA+ cells were TOMATO-negative, indicating that these cells were not derived from postnatal alveolar myofibroblasts (Supplementary Fig. 2C-D).

Given that the transgenic *SMA-CreER* mouse line used in this study enabled lineage labeling even in cells with low levels of *Acta2* transcription (Fig. 2A, control), we investigated whether pre-existing SMA lineage cells reactivated SMA expression during lung regrowth using *SMA-CreER;Ai9* mice. In the control lungs, TOMATO-reporter-positive cells with weak or no SMA expression were observed in the alveolar ducts, and most post-PNX-SMA+ cells were derived from reporter-positive cells (Fig. 2A-B). These findings suggest that post-PNX-SMA+ cells arise from a preexisting SMA-lineage population rather than from *Pdgfra*-lineage alveolar fibroblasts.

**Fig. 2.**
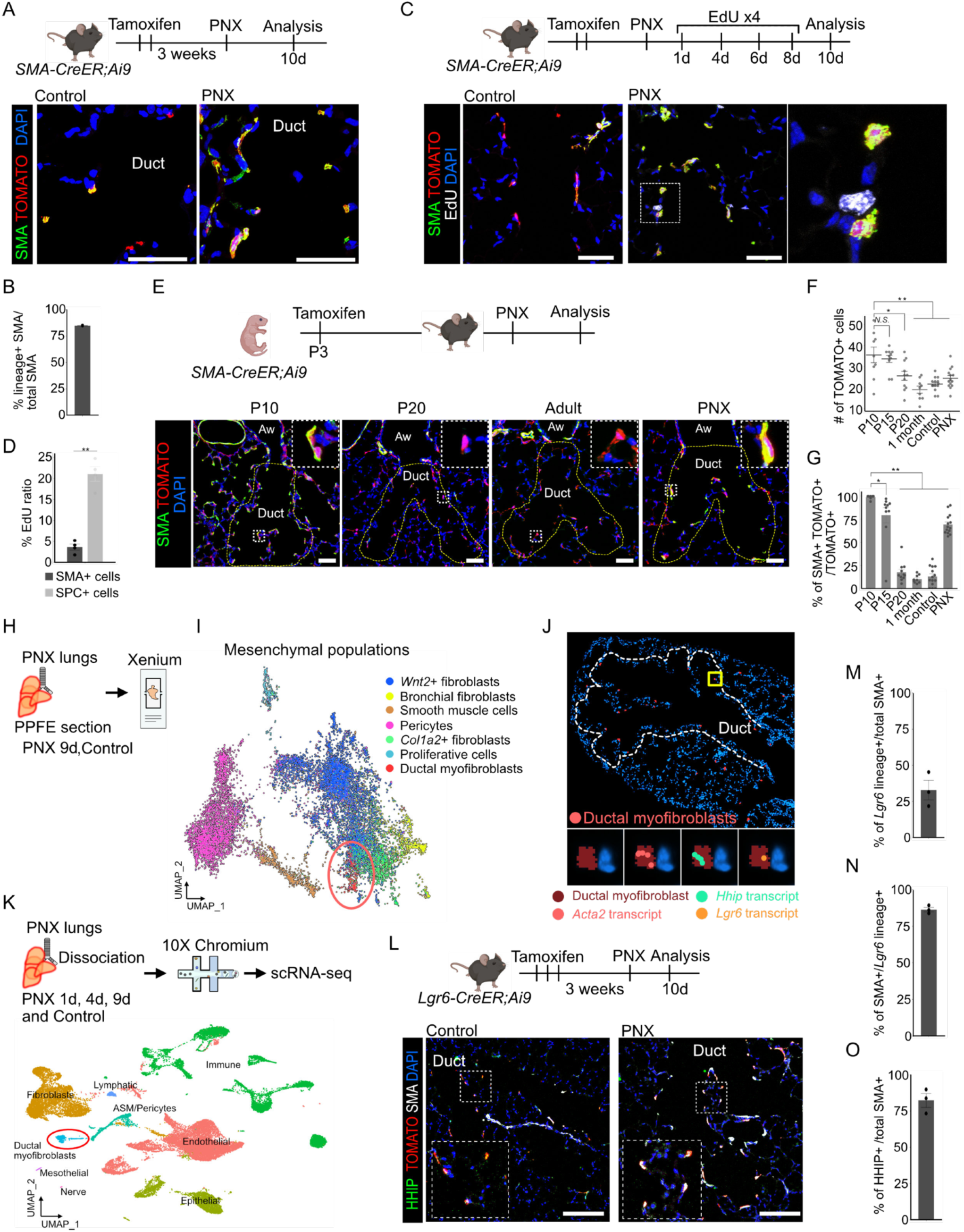
*Lgr6+ Hhip+* ductal myofibroblasts reactivate SMA expression in the regenerating lungs after PNX. A. Schematic diagram of lineage-tracing experiment of SMA+ cells in adult lungs and immunofluorescence images of sections of the alveolar ducts from control (left) and PNX 10d (right) stained for SMA (green) with signal from TOMATO (red). Scale bars, 50 µm. B. Quantification of the ratio of the SMA-lineage+ cells in total SMA+ cells in the alveolar ducts after PNX. (n = 3 mice/group) C. Schematic diagram of EdU incorporation assay in SMA-CreER;Ai9 mice and immunofluorescence images of sections of the alveolar ducts from control (left) and PNX 10d (right) stained for SMA (green) and EdU (white) with signal from TOMATO (red). High-magnification images of the insets drawn by white square. Scale bars, 50 µm. D. Quantification of the ratio of the EdU+ cells in the indicated cell populations at PNX 10d. (n = 4 mice/group) E. Schematic diagram of lineage-tracing experiment of SMA+ cells from neonate and immunofluorescence images of alveolar duct regions at indicated time points stained for SMA (green) with signal from TOMATO (red). High-magnification images of the insets drawn by white squares. Scale bars, 50 µm. F. Quantification of the number of TOMATO+ cells in the alveolar ducts. Each dot indicates the values from each duct. G. Quantification of the ratio of SMA+ TOMATO+ cells in total TOMATO+ cells. Each dot indicates the values from each duct. (n = 3 mice/group) H. Schematics of experiment for Xenium spatial transcriptomic analysis. I. UMAP plot showing mesenchymal population from Xenium analysis. Red circle indicates ductal myofibroblast cluster. J. Xenium spatial plot indicating ductal myofibroblasts. High-magnification of the inset drawn by yellow square shows the detection of indicated transcripts (bottom). K. Schematic diagram of experimental design for scRNA-seq of PNX lungs and UMAP visualization of major cell types of lungs after integration of the all time points. The red circle shows the ductal myofibroblast cluster. L. Schematic diagram of lineage-tracing experiment of *Lgr6*+ cells and immunofluorescence images of lung sections from control (top) and PNX 12d (bottom) stained for HHIP (green) and SMA (white) with signal from TOMATO (red). High-magnification images of the insets drawn by white squares. Scale bars, 150 µm. M. Quantification of the ratio of *Lgr6*-lineage+ cells in total SMA+ cells in the ducts after PNX. (n = 3 mice/group) N. Quantification of the ratio of SMA+ cells in *Lgr6*-lineage+ cells in the ducts after PNX. (n = 3 mice/group) O. Quantification of the ratio of HHIP+ SMA+ cells in total SMA+ cells in the ducts after PNX. (n = 3 mice/group) All data represent the mean ± SEM obtained from at least three independent mice. \**P*<0.05, \*\**P*<0.01.

Next, we assessed proliferative activity in alveoli using an EdU incorporation assay (Fig. 2C). While more than 20% of SFTPC+ AT2 cells, alveolar stem cells, were proliferating after PNX, only minimal EdU incorporation was observed in ductal SMA+ cells (Fig. 2D and Supplementary Fig. 2E). This observation suggests that post-PNX-SMA+ cells do not undergo active proliferation, but may already reside in this location under homeostatic conditions.

Subsequently, we performed lineage tracing using *SMA-CreER;Ai9* mice from neonates to adulthood to further confirm that the post-PNX-SMA+ cells emerging during regeneration were derived from preexisting SMA-lineage cells. By combining lineage tracing with SMA immunostaining, we tracked the dynamics of SMA expression from postnatal development to adult lung regrowth. Tamoxifen was administered at P3 and labeled cells were examined from P10 to adulthood (Fig. 2E-G, Supplementary Fig. 2F-H). The TOMATO reporter-positive cells in the alveolar ducts were maintained over time despite the downregulation of SMA after P20 (Fig. 2E (insets), F, G). In contrast, in the distal alveoli, SMA expression and the number of TOMATO reporter-positive cells progressively declined, with the majority of them disappearing over time (Supplementary Fig. 2F-H). Only a small number of TOMATO reporter- positive cells remained within adult alveoli, including pericytes, airway and vascular smooth muscle. Notably, these persistent TOMATO+ cells showed upregulated SMA expression within 10 days of PNX. Collectively, our findings demonstrated that post-PNX-SMA+ cells derived from preexisting SMA-lineage cells, that are derived neither from adult alveolar fibroblasts nor from transient neonatal distal alveolar myofibroblasts.

We then performed a single-cell-resolution spatial transcriptomic analysis using the 10x Genomics Xenium prime 5K platform to further characterize the reactivated post-PNX-SMA+ cells during regeneration (Fig. 2H). Cluster analysis identified all the major lung cell types, and *Acta2* transcripts were detected in the alveolar ducts (Supplementary Fig. 3A-B). Sub-clustering of the mesenchymal populations further resolved the seven distinct clusters (Fig. 2I). Among them, we identified a specific mesenchymal population enriched in established ductal myofibroblast markers, including *Lgr6*, *Hhip*, and *Cdh4* (Fig. 2I, Supplementary Fig. 3C-D)^15–17^. Consistent with this observation, *Hhip* and *Lgr6* transcripts were detected in the alveolar duct (Supplementary Fig. 3E-F), and spatial mapping confirmed that the cells within the ductal myofibroblast cluster were localized to the alveolar ducts (Fig. 2J). Additionally, a comparable ductal myofibroblast population was independently identified using droplet-based single-cell RNA sequencing (Chromium scRNA-seq) (Fig. 2K). Although ductal myofibroblasts were anatomically contiguous with airway smooth muscle, they lacked the expression of canonical airway smooth muscle markers such as *Mcam* and *Itga8*^36^, indicating that they represent a distinct mesenchymal population (Supplementary Fig. 3G-I).

Subsequently, we performed lineage-tracing analysis using the *Lgr6-CreER;Ai9* mouse line to determine whether *Lgr6*+ ductal myofibroblasts reexpressed SMA *in vivo* after PNX (Fig. 2L, Supplementary Fig. 3J). In control lungs, HHIP+ TOMATO+ (*Lgr6*-linegae) cells were present in the ducts but did not express SMA. Following PNX, 32.8% ± 6.8% of post-PNX-SMA+ cells were derived from the *Lgr6* lineage, and 86.3% ± 1.6% of *Lgr6*-lineage cells were positive for SMA (Fig. 2M-N). Additionally, 82.4% ± 4.8% of SMA+ cells in the duct were HHIP+ (Fig. 2O), indicating *Lgr6*+ ductal myofibroblasts reactivate SMA during lung regrowth and become post-PNX-SMA+ cells. Three-dimensional analysis further revealed that these *Lgr6*-lineage cells exhibited spindle-shaped morphology, distinct from alveolar fibroblasts and pericytes, even under homeostatic conditions (Supplementary Fig. 3J). These findings demonstrate that postnatal *Lgr6*+ ductal myofibroblasts serve as the origin of post-PNX-SMA+ cells and that these two populations represent identical cell types. Following PNX treatment, these cells undergo functional SMA reactivation, which contributes to compensatory lung regeneration. Based on these distinct features, we designated these cells as PNX-activated ductal myofibroblasts (PA-DMFs).

### Autocrine myogenic induction in PA-DMFs reactivate contractile program

We explored the mechanism driving acquisition of a contractile program in PA-DMFs following PNX by conducting sub-clustering analysis of our scRNA-seq dataset. The ductal-myofibroblast/PA-DMF population was subdivided into four distinct clusters to define the unique molecular features of these myofibroblasts (Fig. 3A-B). The proportion of cluster 2 increased progressively after PNX, suggesting that PA-DMFs were molecularly distinguishable from ductal myofibroblasts (Fig. 3B-D). Consistent with this observation, *Acta2* expression was elevated in cluster 2, as corroborated by our *in vivo* observations (Fig. 3E). Cluster-specific marker analysis and Reactome pathway enrichment revealed that PA-DMFs were significantly enriched for terms related to collagen production and ECM organization (Supplementary Fig. 3K). Additionally, several genes associated with myogenic potential, including *Col15a1, Ltbp2, Ccn4,* and *Fndc1*, were markedly upregulated in PA-DMFs (Fig. 3F). Among these, *Ccn4* encodes a secreted factor known to regulate cell reactivation and state transitions. We therefore intranasally administered recombinant CCN4 to mice prior to PNX. Notably, the intranasal administration of recombinant CCN4 alone elicited SMA expression in ductal myofibroblasts without PNX, suggesting that CCN4 contributes to the activation of this population. (Fig. 3G). Furthermore, pathways related to focal adhesion and regulation of the actin cytoskeleton were significantly upregulated (Supplementary Fig. 3L), suggesting enhanced contractile activity within PA-DMFs. Additionally, immunofluorescence analysis consistently showed a significant increase in phosphorylated myosin light chain (pMLC)+ SMA+ cells in ductal myofibroblasts after PNX (Fig. 3H-I). In contrast, intraperitoneal administration of ML-7, an MLCK inhibitor, decreased MLC phosphorylation in ductal myofibroblasts after PNX, resulting in enlarged alveolar ducts, suggesting the importance of contractile force in the maintenance of alveolar architecture (Fig. 3J-L). Together, these findings determine a mechanism of alveolar tissue stabilization post-PNX, where myofibroblasts secrete myogenic factors, such as CCN4, to reactivate a myogenic program that converts them into contractile PA-DMFs to preserve alveolar architecture.

**Fig. 3.**
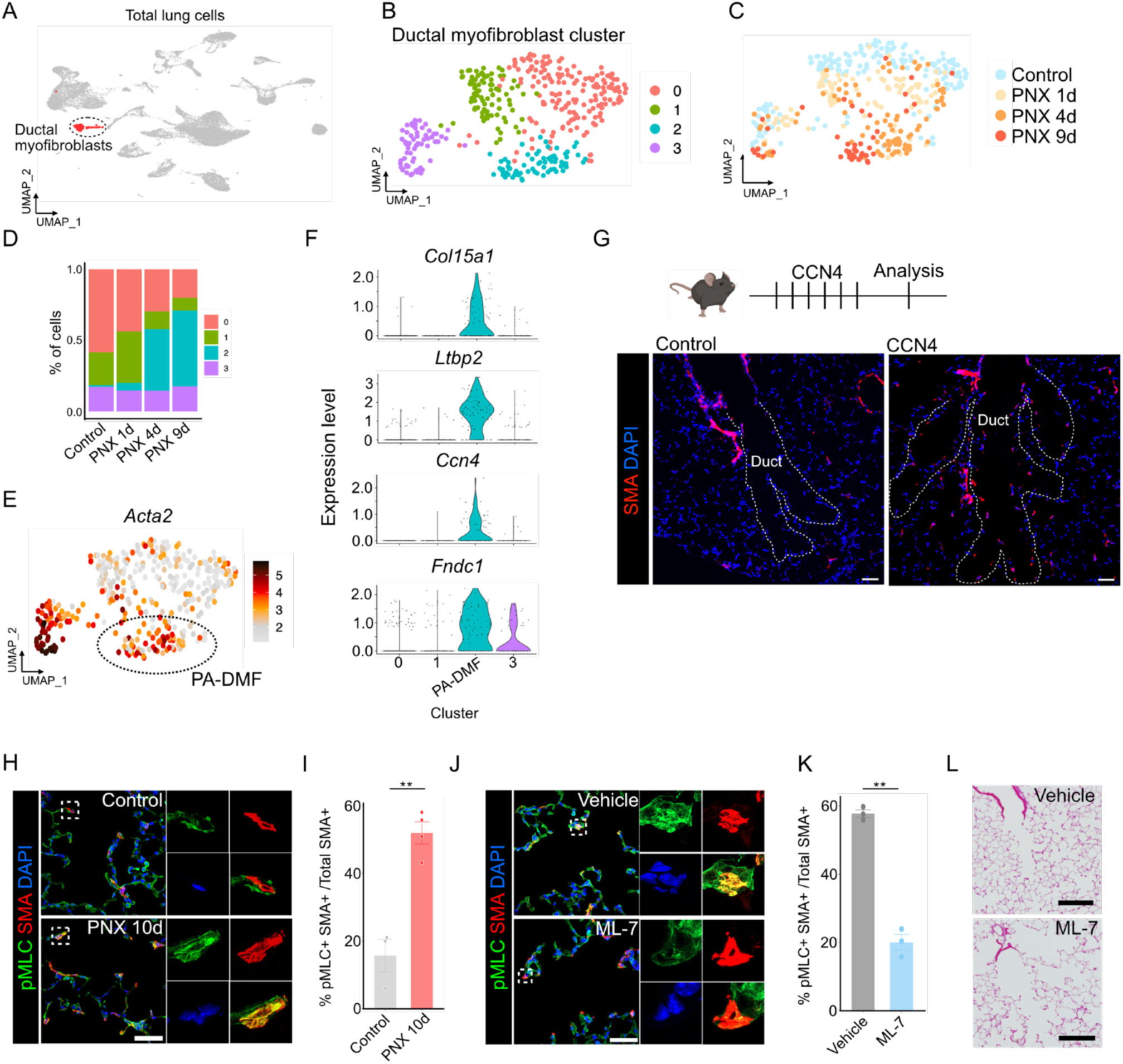
Ductal myofibroblasts shift to PA-DMFs via autocrine CCN4 to reactivate contractile program and maintain alveolar architecture after PNX. A. UMAP plot highlighting ductal myofibroblast cluster (red) in whole lung cells. B. UMAP plot from sub-clustering analysis of the ductal myofibroblast cluster. C. UMAP plot showing temporal transition of the ductal myofibroblasts after PNX. D. Bar graph showing the ratio of each cluster at each time point. E. UMAP plot showing the expression level of *Acta2* gene. F. Violin plots indicating the expression of indicated genes. G. Immunofluorescence images of lung sections from control (top) and CCN4-treated mice (bottom) stained for SMA (red). Scale bars, 50 µm. H. Immunofluorescence images of lung sections from control (top) and PNX 10d (bottom) stained for pMLC (green) and SMA (red). High-magnification images of the insets drawn by white squares (right). Scale bars 50 µm. I. Quantification of the ratio of pMLC+ SMA+ cells in total SMA+ cells in the alveolar ducts. (n = 3-4 mice/group) J. Immunofluorescence images of lung sections from control (top) and ML-7 treated mice (bottom) at PNX 10d stained for pMLC (green) and SMA (red). High-magnification images of the insets drawn by white squares (right). Scale bars 50 µm. K. Quantification of the ratio of pMLC+ SMA+ cells in total SMA+ cells in the alveolar ducts. (n = 3 mice/group) L. H&E staining of lung sections from control (top) and ML-7 treated mice (bottom) at PNX 10d. Scale bars, 200 µm. All data represent the mean ± SEM obtained from at least three independent mice. \*\**P*<0.01.

### Human *LGR6*+ fibromyocytes are homologous to mouse *Lgr6*+ ductal myofibroblasts

Recent advances in single-cell transcriptomics have uncovered the cellular landscape of the human lung^37–41^; however, the diversity and functional roles of mesenchymal populations still remain unclear. Therefore, we investigated whether the ductal myofibroblasts identified in mice were conserved in human lungs by analyzing publicly available scRNA-seq data from human distal airways (Fig. 4A, Supplementary Fig. 4A)^38^. Sub-cluster analysis of the mesenchymal population revealed that markers of mouse ductal myofibroblasts, including *LGR6*, *HHIP*, and *ASPN*, were significantly enriched within the fibromyocyte cluster (Fig. 4A-B). Reactome pathway enrichment analysis further indicated that fibromyocytes exhibited gene signatures associated with muscle contraction similar to those of mouse ductal myofibroblasts (Fig. 4C). Next, we extracted 294 fibromyocyte-specific marker genes, converted them into mouse orthologs, and projected them onto a mouse lung scRNA-seq dataset to compare the molecular similarities between human fibromyocytes and mouse ductal myofibroblasts (Fig. 4D). The average expression of these fibromyocyte markers was strongly enriched in ductal myofibroblast clusters in the mouse lung dataset, in addition to airway smooth muscle (Fig. 4D), supporting a close molecular relationship between the two populations. Among the various markers analyzed, we selected *SBSPON* and *SYNPO2* as representative markers of human fibromyocytes and examined their expression in mouse lungs to further validate our finding. Analysis of mouse scRNA-seq data showed that *Sbspon* and *Synpo2* were highly expressed in ductal myofibroblast clusters (Supplementary Fig. 4B-C). Additionally, *in situ* hybridization confirmed that transcripts of *Sbspon* and *Synpo2* were detected in *Lgr6*-lineage positive ductal myofibroblasts in mouse lungs (Supplementary Fig. 4D-E), suggesting that human fibromyocytes and mouse ductal myofibroblasts have conserved molecular characteristics. Subsequently, we evaluated the anatomical localization of fibromyocytes in human lungs. Based on scRNA-seq analysis, fibromyocytes were predicted to reside in distal airway regions, including the terminal respiratory bronchioles (TRB), which form the interface between the airway and alveolar compartments (Fig. 4E)^42^. Consistent with this prediction, we identified SCGB3A2+ SFTPB+ TRB-specific epithelial cells^38,39^ and intermittent clusters of SMA+ cells beneath the TRB epithelium (Fig. 4F). *In situ* hybridization further revealed that *LGR6+ ACTA2+* fibromyocytes constituted 73.8% ± 4.0% of *ACTA2*+ cells within the TRB region (Fig. 4G-H), whereas such cells were not detected in the alveoli (Supplementary Fig. 4F). Similar results were obtained for four independent normal human lungs from different donors (Supplementary Fig. 5). Notably, mouse ductal myofibroblasts are localized in the alveolar ducts, which represent the transitional regions connecting the airway and alveolar compartments. Thus, human fibromyocytes and mouse ductal myofibroblasts exhibit striking similarities in their molecular identity and anatomical localization. Taken together, these findings demonstrate that *Lgr6*+ ductal myofibroblasts are evolutionarily conserved in the human lungs as *LGR6*+ fibromyocytes, sharing molecular signatures and anatomical distribution.

**Fig. 4.**
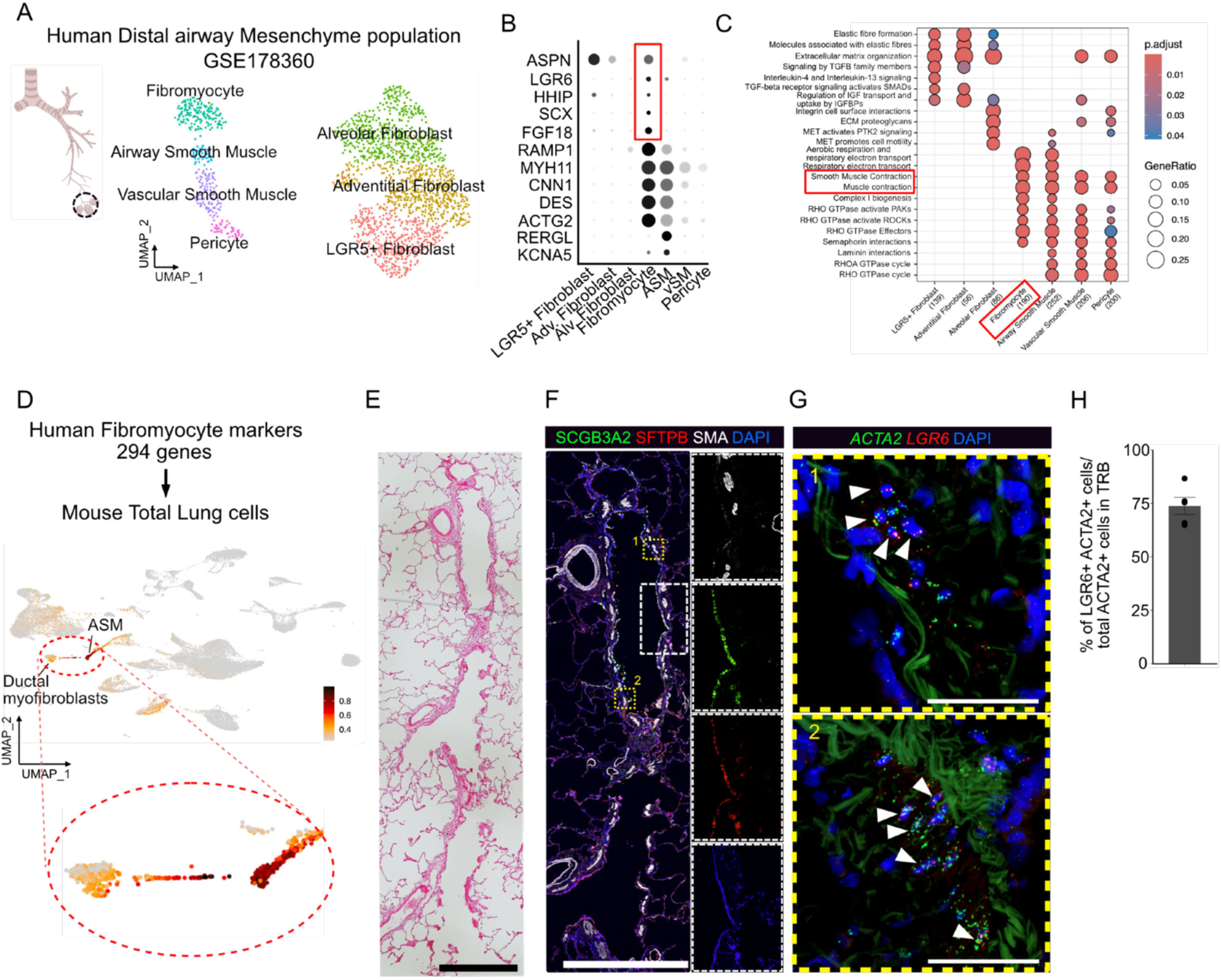
Cross-species scRNA-seq analysis reveals that mouse ductal myofibroblasts were anatomically and molecularly conserved in human fibromyocytes. A. Schematic diagram of human airway tree and UMAP visualization of mesenchyme population in human distal airway from GSE178360. B. Dot plot showing cluster-specific marker genes corresponding to the annotation in (A). The red rectangle indicates fibromyocyte-specific genes. C. Reactome pathways enriched in each cluster. The red rectangle indicates terms related to contractile forces. D. UMAP plot showing the average expression of human fibromyocyte-specific marker genes in mouse total lung cells. E. H&E staining of the terminal airway in human lung tissue. Scale bar, 1 mm. F. Immunofluorescence image of human terminal airway stained for SCGB3A2 (green), SFTPB (red), and SMA (white). High-magnification single-color images of the insets drawn by white squares. Scale bar, 1 mm. G. Detection of transcripts of ACTA2 (green) and LGR6 (red) in the region drawn by yellow circles in (E). White arrowheads indicate double positive cells. Scale bars, 50 µm. H. Quantification of the ratio of LGR6+ ACTA2+ cells in total ACTA2+ cells in the terminal airways.

### Developmental trajectory of PA-DMFs from embryonic distal-most SMA⁺ cells

Next, we focused on the developmental process through which ductal SMA⁺ cells appear at the distal-most airways during lung morphogenesis to better understand how cell fate is determined (Fig. 5A, white circles). Whole-mount imaging at the late pseudoglandular stage (E14.5, E15.5) revealed that SMA+ cells were present around SOX2+ airway epithelial cells, consistent with airway smooth muscle cells, and in the more distal SOX2 negative regions corresponding to future alveolar domains (Fig. 5B)^43^. At the canalicular (E16.5) stage, SMA+ cells at these distal tips became less densely connected compared to the proximal airway smooth muscle, forming discontinuous bundles reminiscent of ductal myofibroblast distribution (Fig. 1I-J, and Fig. 5B). Subsequently, we examined the expression of MCAM, a canonical airway smooth muscle marker to determine whether these distal SMA+ cells correspond to ductal myofibroblasts^36^. Because scRNA-seq and immunostaining confirmed that ductal myofibroblasts did not express MCAM at either the gene or protein levels (Supplementary Fig. 3G-H), MCAM served as a marker to distinguish mature airway smooth muscle from ductal myofibroblasts, although endothelial cells were also MCAM-positive. Consistent with our results for the PNX lungs (Supplementary Fig. 3H), MCAM expression in the embryonic lungs was restricted to the airway smooth muscle in the subepithelial compartment (Fig. 5C, yellow dashed lines). In contrast, the SMA+ cells located distal to the SOX2+ airway regions were MCAM-negative (Fig. 5C; white brackets). These findings indicate that the embryonic SMA+ population can be subdivided into two distinct groups: MCAM+ airway smooth muscle surrounding the SOX2+ airway epithelium, and MCAM-distal SMA+ cells located in the future alveolar region distal to the SOX2+ airway epithelium (Fig. 5D).

**Fig. 5.**
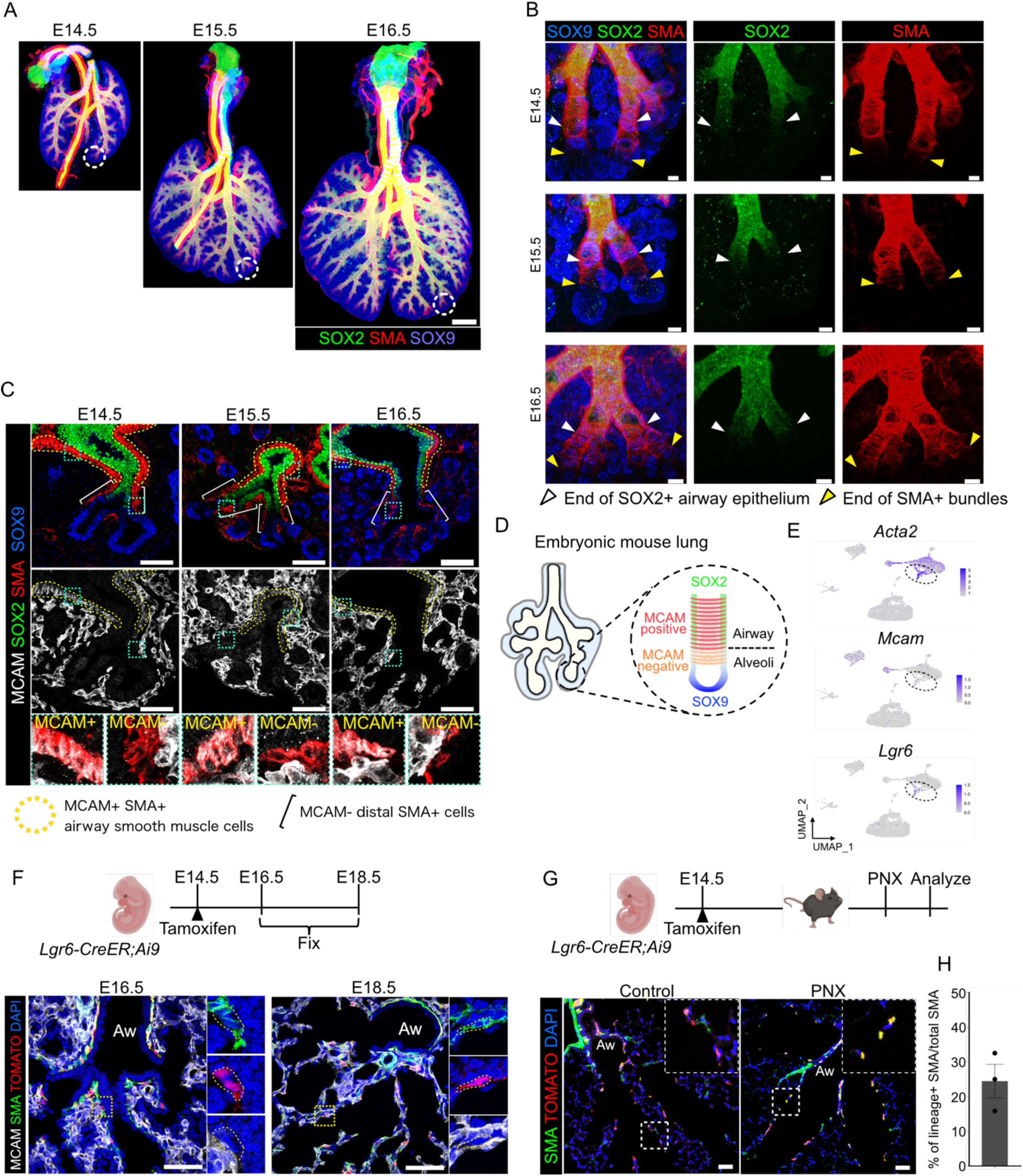
MCAM- SMA+ cells located at the distal end of the embryonic lungs contribute to PA-DMF in adult lungs. A. Whole-mount images of embryonic lungs stained for SOX2 (green), SMA (red), and SOX9 (blue). Scale bars, 500 µm. B. Three-dimensional images of cleared embryonic lungs stained for SOX9 (blue), SOX2 (green), and SMA (red) at the indicated embryonic days. White and yellow arrowheads indicate the end of SOX2+ airway epithelium and the end of SMA+ bundles in the airway branches, respectively. Scale bars, 30 µm. C. Immunofluorescence images of embryonic lung sections stained for MCAM (white), SOX2 (green), SMA (red), and SOX9 (blue). Yellow dashed lines and white brackets indicate MCAM+ SMA+ airway smooth muscles and MCAM- distal-most SMA+ cells, respectively. The bottom panels show MCAM+ SMA+ (left in each time point) and MCAM- SMA+ (right in each time point) cells. Scale bars, 50 µm. D. Schematic diagram of the distribution of MCAM- and MCAM+ SMA+ cells in the embryonic lungs. E. Feature plots of E15.5 lung dataset showing the expression of the indicated genes. F. Schematic diagram of lineage-tracing experiment of *Lgr6*+ cells in the embryonic lungs and immunofluorescence images of lung sections stained for MCAM (white) and SMA (green) with signal from TOMATO (red) at indicated time points. Yellow dashed lines indicate lineage positive MCAM- SMA+ cells in the alveoli. Scale bars, 50 µm. G. Schematic diagram of lineage-tracing experiment of *Lgr6*+ cells from embryo to adult and immunofluorescence images of lung sections from control (left) and PNX 12d (right) stained for SMA (green) with signal from TOMATO (red). High-magnification images of the insets drawn by white squares. Scale bars 50 µm. H. Quantification of the ratio of *Lgr6*+ lineage positive SMA+ cells in total SMA+ cells in the alveolar ducts. (n = 3 mice/group) Data represent the mean ± SEM obtained from three independent mice.

Thereafter, we investigated whether these MCAM- distal SMA+ cells gave rise to ductal myofibroblasts. scRNA-seq analysis of E15.5 lung^6^ showed that *Mcam- Acta2+* cluster expressed *Lgr6*, a marker of ductal myofibroblasts in adult lungs (Fig. 5E). Lineage-tracing analysis of *Lgr6*+ cells from embryonic stages to adulthood was then performed. Tamoxifen was administered to pregnant *Lgr6- CreER;Ai9* dams at E14.5. TOMATO+ cells were subsequently detected in a subset of distal SMA+ cells in the alveoli of embryonic lungs from E16.5–E18.5, in addition to airway smooth muscle cells, confirming that *Lgr6* lineage labeling marks MCAM- distal SMA+ cells (Fig. 5F). In adult lungs, TOMATO+ cells labeled at E14.5, were observed within the alveolar ducts (Fig. 5G). Although these cells lacked SMA expression under homeostatic conditions, they re-expressed SMA after PNX, which was consistent with the behavior of PA-DMFs (Fig. 5G-H). These results indicated that embryonic *Lgr6*-lineage MCAM- distal SMA+ cells give rise to ductal myofibroblasts during the perinatal period and reactivate SMA expression following PNX in adulthood. This suggests that embryonically established ductal myofibroblasts contribute to a stable and lifelong population that maintains alveolar architecture and is necessary for both alveolar development and regeneration.

### Common origin of ductal myofibroblasts and distal airway smooth muscle

Because *Lgr6-CreER;Ai9* mice were labeled with a subset of airway smooth muscle cells in addition to ductal myofibroblasts (Fig. 5F), we hypothesized that MCAM- distal SMA+ cells serve as common progenitors for both lineages. Accordingly, we assessed the proliferative activity of MCAM- distal SMA+ cells using the EdU incorporation assay to evaluate progenitor potential. EdU was injected to pregnant dams at each embryonic stages from E14.5 to E17.5, and the embryonic lungs were analyzed 4 h later (Supplementary Fig. 6A). At E14.5, 27.5% ± 1.8% of MCAM- distal SMA+ cells incorporated EdU, whereas proliferation was significantly lower in proximal SMA+ cells surrounding large bronchi (Supplementary Fig. 6B-D, yellow arrowheads). While proximal airway smooth muscle proliferation remained below 5% throughout the stages examined, MCAM- SMA+ cells exhibited 17.0% ± 1.6% at E15.5 before declining after E16.5. These results indicated that MCAM- SMA+ cells are highly proliferative, with activity both spatially and temporally restricted to the pseudoglandular stage.

Subsequently, we performed EdU pulse-chase experiments initiated at E14.5 to determine whether these proliferating distal SMA+ cells contribute to airway smooth muscle (Supplementary Fig. 6E). At E15.5, EdU+ SMA+ cells were identified at the distal ends and within the MCAM+ airway smooth muscles (Supplementary Fig. 6F-G). Consistent with previous reports^44,45^, these labeled cells remained present in the airway smooth muscle through P0, demonstrating that the MCAM- distal SMA+ cells migrated and contributed to airway smooth muscle formation (Supplementary Fig. 6H). Collectively, these findings established that MCAM- distal SMA+ cells are the common origin of both ductal myofibroblasts and airway smooth muscle cells during embryonic lung development.

### TGF-β signaling activates SMA+ cell differentiation from the embryonic progenitors

To investigate the molecular mechanisms initiating the differentiation of ductal myofibroblasts and distal airway smooth muscle cells, we analyzed an scRNA-seq dataset from E15.5 lung^6^ to identify epithelial factors involved in epithelial-mesenchymal interactions. Sub-clustering analysis identified three epithelial populations: *Sox2*+ airway epithelium, *Sox9*+ distal bud epithelium, and *Hopx*+ stalk epithelium located between the airway and distal bud regions (Fig. 6A). Given that distal SMA+ cells emerged adjacent to the *Hopx*+ stalk region, but not the *Sox9*+ distal buds, we searched for signaling molecules enriched in the stalk epithelium that could regulate SMA+ cell differentiation. Among the growth factors associated with lung development, *Tgfb2* expression was markedly enriched in stalk epithelial cells (Fig. 6A-B). *In situ* hybridization confirmed the presence of *Tgfb2* transcripts in epithelial cells located above *Sftpc+* lung buds (Fig. 6C). Therefore, we examined whether subepithelial SMA+ cells respond to TGF-β ligands. *In situ* hybridization detected the expression of *Tgfbi*, a downstream target gene of TGF-β signaling^46^, in subepithelial mesenchyme above *Sftpc*+ lung buds (Supplementary Fig. 6I). The *Tgfbi*+ cells also expressed *Acta2* (Fig. 6D). In addition, phosphorylated SMAD2/3, key downstream mediators of the TGF- β signal cascade, were detected in subepithelial SMA+ cells in embryonic lungs (Fig. 6E), indicating active TGF-β signaling in subepithelial SMA+ cells.

**Fig. 6.**
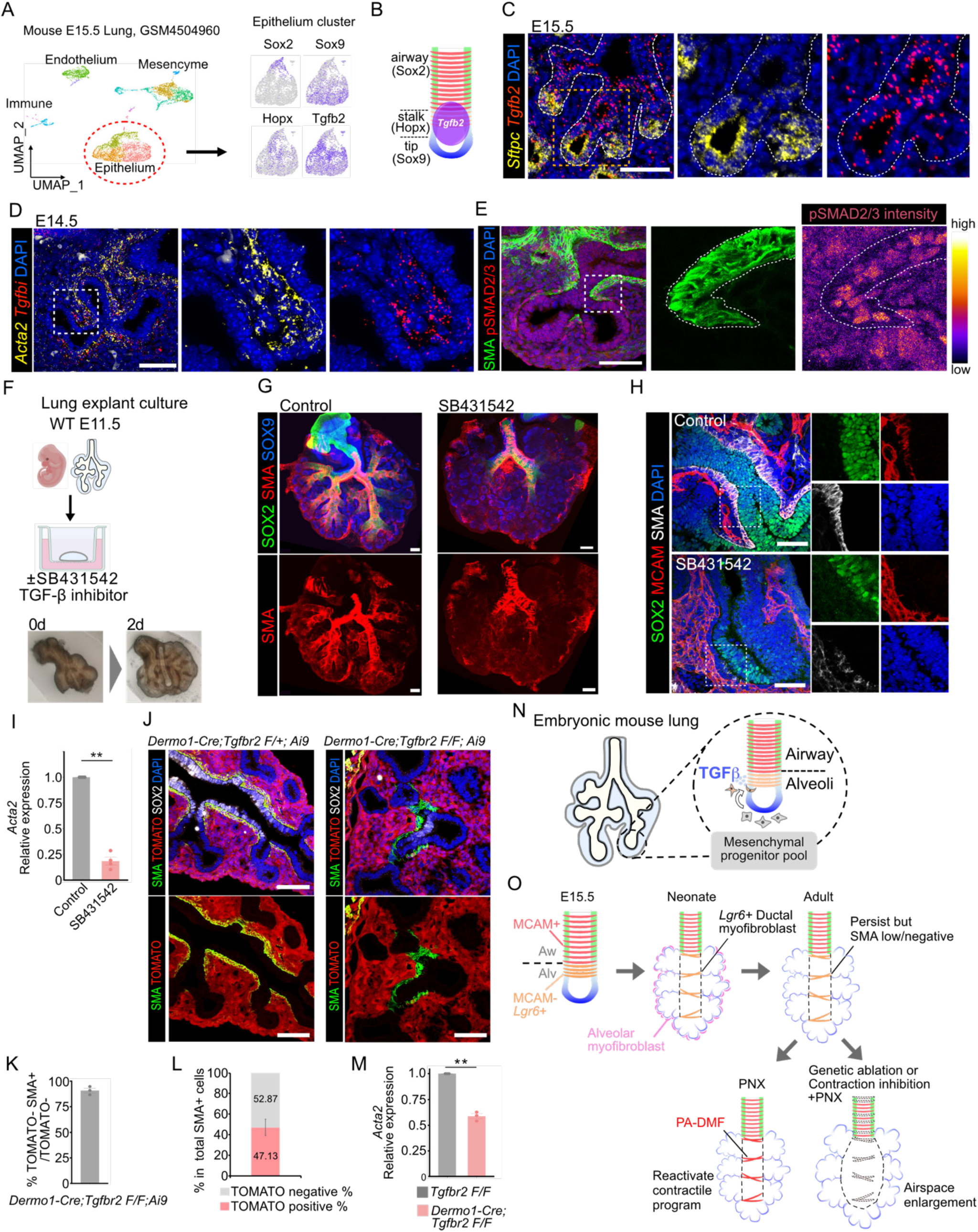
The stalk epithelium-derived TGF-β signal activate the differentiation of SMA+ cells from mesenchymal progenitors. A. UMAP visualization of total lung cells from E15.5 embryo (GSM4504960, left) and feature plots showing the indicated genes in epithelium clusters. B. Schematic diagram of spatial distribution of *Tgfb2* expression in the embryonic lungs inferred from scRNA-seq analysis. C. Detection of transcripts of *Sftpc* (yellow) and *Tgfb2* (red) in embryonic lungs. White dashed lines indicate epithelial layers. High-magnification images of the insets drawn by yellow squares. Scale bars, 50 µm. D. Detection of transcripts of *Acta2* (yellow) and *Tgfbi* (red) in E14.5 lungs. High-magnification images of the insets drawn by white squares. Scale bars, 50 µm. E. Immunofluorescence images of section stained for SMA (green) and pSMAD2/3 (red) at E14.5. High-magnification images of the insets drawn by white squares. White dashed lines indicate epithelial layers. Visualization of the intensity of pSMAD2/3 signals (right). Scale bars, 50 µm. F. Schematic diagram of organ culture experiment and representative images of lung explant before and after culture. G. Whole-mount images of lung explants cultured for 2 d with (right) or without (left) SB431542 stained for SOX2 (green), SMA (red), and SOX9 (blue). Scale bars, 100 µm. H. Immunofluorescence images of sections of lung explants cultured with (bottom) or without (top) SB431542 stained for SOX2 (green), MCAM (red), and SMA (white). Scale bars, 50 µm. I. Comparison of *Acta2* mRNA expression levels in organ culture with or without SB431542. (n = 4 mice/group) J. Immunofluorescence images of lung sections from *Dermo1-Cre;Tgfbr2 F/+;Ai9* (control, left) and *Dermo1-Cre;Tgfbr2 F/F;Ai9* (right) stained for SMA (green) and SOX2 (white) with signal from TOMATO (red). Scale bars, 50 µm. K. Quantification of the ratio of TOMATO- SMA+ cells in the subepithelial TOMATO- cells. (n = 3 mice/group) L. Quantification of the ratio of TOMATO positive and negative cells in total SMA+ cells in lungs Dermo1-Cre;Tgfbr2 F/F;Ai9 mice at E15.5. Data represent the mean from three independent mice. M. Comparison of *Acta2* mRNA expression levels in lungs at E15.5. Data represent the mean ± SEM obtained from three independent mice. (n = 3 mice/group) N. Schematic diagram of SMA+ cell differentiation stimulated by TGF-β signaling derived from the stalk epithelium in embryonic lungs. O. Summary of our findings. MCAM- Lgr6+ distal SMA+ cells in the embryonic lungs give rise to ductal myofibroblasts which are maintained beyond postnatal phase while downregulate SMA expression. These cells reactivate SMA expression and contractile program to stabilize alveolar architecture during lung regeneration. All data represent the mean ± SEM obtained from at least three independent mice. \*\**P*<0.01.

Next, we utilized an embryonic lung explant culture system that recapitulates developmental processes *ex vivo* to test whether TGF-β signaling regulates SMA+ cell differentiation (Fig. 6F). In the control explants, SMA expression was observed along the developing airway branches. In contrast, treatment with SB431542, a TGF-β receptor inhibitor, resulted in complete loss of SMA expression within the lung tissue, although SMA+ cells remained in the extrapulmonary regions (Fig. 6G). Immunofluorescence corroborated these findings (Fig. 6H). Consistently, quantitative PCR analysis demonstrated a marked reduction of *Acta2* mRNA expression in explants treated with TGF-β inhibitor (Fig. 6I).

Because pharmacological inhibition affects multiple cell types in the explant culture system, including epithelial and endothelial cells, we examined the role of TGF-β signaling specifically in mesenchymal cells using *Dermo1-Cre;Tgfbr2 F/F* conditional knockout mice. While TOMATO+ SMA+ cells were readily detected in the subepithelial regions of control lungs, *Dermo1-Cre;Tgfbr2 F/F;Ai9* mutant lungs displayed disrupted SMA bundle formation (Fig. 6J). Although SMA+ cells were still observed in certain regions, many TOMATO- cells give rise to SMA+ cells, possibly reflecting incomplete recombination or compensation from alternative cell sources (Fig. 6J-L). Quantitative PCR further showed that *Acta2* expression was significantly reduced in the lungs of mutant mice (Fig. 6M). Taken together, these results demonstrate that activation of TGF-β signal in subepithelial mesenchyme is required for the differentiation of SMA+ ductal myofibroblasts and distal airway smooth muscle cells during embryonic lung development (Fig. 6N).

## Discussion

In this study, we identified ductal myofibroblasts as a critical mesenchymal subset of cells required for maintaining the alveolar architecture during lung regrowth following PNX. Using lineage tracing, spatial transcriptomics, and single-cell RNA sequencing, we demonstrated that resident *Lgr6*+ *Hhip*+ SMA- ductal myofibroblasts in the alveolar ducts convert into PA-DMFs during post-PNX lung regrowth through autocrine CCN4 signaling, which drives SMA expression and reactivates a contractile program. Functional ablation experiments further demonstrated that PA-DMFs are required to maintain the alveolar structure and avoid an emphysema phenotype following PNX. Together, these findings revealed a previously underappreciated structural cell population that stabilizes the alveolar architecture during lung regrowth (Fig. 6O).

Ductal myofibroblasts were first identified through scRNA-seq analysis of mesenchymal populations in developing lungs^15^. Because SMA has been widely used as a marker of myofibroblasts and is expressed in both alveolar and ductal myofibroblasts^5^, these populations have not been clearly distinguished in earlier studies. Subsequent analyses showed that ductal myofibroblasts are characterized by the expression of markers, such as *Hhip*, *Lgr6*, and *Cdh4*, and lineage-tracing studies demonstrated that they are still detectable in the lungs after the completion of postnatal alveologenesis, in contrast to alveolar myofibroblasts^15–17^. A recent study also identified ductal myofibroblast-like populations by analyzing Hippo signaling during alveolar morphogenesis. In this study, the dynamic regulation of Yap/Taz activity in *Acta2*-lineage cells controlled contractile gene expression, and the disruption of this pathway led to the accumulation of aberrant cells expressing ductal myofibroblast markers^16^. Similar populations have been observed in models in which apoptosis of alveolar myofibroblasts is inhibited^17^. Although these studies suggest the presence of ductal myofibroblasts, their functional roles remain unclear. Our findings extend these observations by demonstrating that ductal myofibroblasts function as a long-lived quiescent mesenchymal population that can be reactivated to exert contractile forces during lung regrowth. Genetic ablation of SMA+ cells and pharmacological inhibition of the contractile ability resulted in regional emphysematous changes following PNX, likely reflecting the unique spatial distribution of ductal myofibroblasts in the alveolar compartment. Nevertheless, these findings support the role of these cells in maintaining the alveolar architecture. Together, these findings suggest that ductal myofibroblasts represent a resident mesenchymal population that is preserved after development and is reengaged to maintain tissue architecture during regeneration, highlighting the general mechanism by which structural cell populations support tissue integrity in adults.

Our scRNA-seq analysis revealed that ductal myofibroblasts upregulate myogenic factors following PNX. Among these, CCN4 has emerged as a candidate driving activation of PA-DMFs from SMA- ductal myofibroblasts, as exogenous CCN4 administration is sufficient to induce SMA expression in even the absence of PNX. Furthermore, given that MLC phosphorylation is essential for maintaining alveolar architecture, these findings suggest that CCN4-mediated PA-DMF induction serves as a central mechanism reactivating the contractile forces required for structural stability of the post-PNX alveoli.

CCN4 is a member of the CCN family of matricellular proteins and has been implicated in a wide range of cellular processes, including cell adhesion, proliferation, and differentiation, as well as inflammation and tissue repair^47^. In skeletal muscles, CCN4 secreted by fibro-adipogenic progenitors supports muscle stem cell proliferation and commitment, and its decline with age contributes to impaired regenerative capacity, which can be restored by exogenous CCN4^48^. In addition to CCN4, other factors upregulated in PA-DMFs are also linked to muscle-associated functions. *Col15a1* deficiency results in myopathic phenotypes and structural instability in skeletal and cardiac muscle^49^, whereas FNDC1 promotes myoblast differentiation and has been associated with age-related decline in muscle function^50^. Together, these observations support the notion that ductal myofibroblasts acquire a muscle-like functional program (e.g., generation of contractile forces) upon activation. The temporal induction of these factors, particularly on day 9 after PNX, suggests that the initial induction of PA-DMFs from SMA- ductal myofibroblasts is likely driven by mechanical cues such as rapid expansion of the remaining lung lobes during regrowth^24^. Subsequently, these cells may reinforce and sustain their activated state through autocrine signaling mediated by factors such as CCN4. Although CCN4 has been implicated in fibrotic responses^51^, our findings raise the possibility that controlled activation of ductal myofibroblasts through myogenic factors promote the stabilization of the alveolar architecture.

Fibromyocytes, the human mesenchymal counterparts of mouse ductal myofibroblasts, were found to localize to the TRB in human lungs. The TRB represents a specialized anatomical compartment connecting the conducting airways and alveoli and is considered a human-specific structure absent in the mouse lung^38,39,42^. Recent studies have highlighted this region as a niche for SCGB3A2+ epithelial cells (also referred to as AT0^38^ or respiratory airway secretory cells^39^), which contribute to alveolar epithelial lineages. Despite its potential importance, the TRB remains poorly characterized owing to its small size and limited accessibility^52^. Emerging evidence suggests that the TRB region is a key site of early pathological changes in COPD^53,54^. High-resolution micro-CT analyses have shown that in severe COPD, the terminal bronchioles are reduced in number and exhibit increased wall thickness, decreased lumen area, and loss of alveolar attachments^55,56^. Notably, these structural alterations precede the development of emphysematous changes in the alveoli^57^, raising the possibility that local mesenchymal populations contribute to early remodeling of this region. Fibromyocytes may represent a candidate structural cell type involved in these changes. Furthermore, integration of lung function traits and COPD-associated GWAS studies have identified smooth muscle cells as a major cell type linked to disease susceptibility^58^. *HHIP*, a fibromyocyte marker, is a well-established COPD susceptibility gene^59–61^. In addition to its role as a regulator of Hedgehog signaling, HHIP has also been implicated in oxidative stress responses^62^. Therefore, dysregulation of HHIP-dependent pathways in fibromyocytes could impair contractile function and compromise local structural integrity, thereby contributing to early remodeling of the TRB. Our findings further suggested that ductal myofibroblasts acquire contractile properties during development and persist into adulthood. This raises the possibility that early life insults such as mechanical ventilation may induce long-lasting alterations in these cells. Such a mechanism may help explain why bronchopulmonary dysplasia is associated with an increased risk of COPD later in life^63–65^. In addition to COPD, idiopathic pulmonary fibrosis is also associated with structural abnormalities in the TRB region but with distinct morphological features, including airway wall thickening accompanied by dilation of the airway lumen^66^. These observations suggest that the TRB is a common site of early structural alteration across major pulmonary diseases, albeit with disease-specific remodeling patterns. However, because small airways contribute less than 10% of total airway resistance, pathological changes in this region are often not detected by standard pulmonary function tests, posing a challenge for early diagnosis^67,68^. However, the role of fibromyocytes in this disease remains unclear. These cells may contribute to airway obstruction through hyperplasia or increased contractility; alternatively, their dysfunction or loss, potentially induced by environmental stressors such as cigarette smoke, may promote emphysematous changes. Although these possibilities require further investigation, the identification of a mouse counterpart of human TRB-associated fibromyocytes provides a framework for experimentally dissecting their roles in disease pathogenesis.

In summary, we identified ductal myofibroblasts as a mesenchymal population that plays a role in maintaining the alveolar architecture by shifting to PA-DMFs upon lung regrowth. These cells are established during development, persist as quiescent populations in adulthood, and are reactivated to exert contractile functions in response to regenerative cues. Our findings further demonstrate that this population shows molecular and anatomical conservation in the human lungs as *LGR6*+ fibromyocytes. Together, these results support a model in which developmentally established mesenchymal cells are retained to maintain the tissue architecture in the adult lung.

## Methods

### Mice

All mice were bred and housed in a specific pathogen-free mouse facility at constant temperature (18- 23 °C) and humidity (40-60%) in sterilized plastic cages. A 12h-light/12h-dark cycle was used. *Pdgfra^tm1.1(cre/ERT2)Blh/J^*(Pdgfra-CreER)^69^ (stock number 032770, Jackson laboratory), Tg(Acta2- cre/ERT2)12Pcn (SMA-CreER)^70^, B6.129P2-*Lgr6^tm2.1(cre/ERT2)Cle^*^/J^ (Lgr6-CreER)^71^ (stock number 016934, Jackson laboratory), B6.Cg-*Gt(ROSA)26Sor^tm9(CAG-tdTomato)Hze^*^/J^ (Ai9)^72^ (stock number 007909, Jackson laboratory), B6;129-*Tgfbr2^tm1Karl^*^/J^ (Tgfbr2flox)^73^ (stock number 012603, Jackson laboratroy), B6.129X1- *Twist2^tm1.1(cre)Dor^*^/J^ (Dermo1-Cre)^74^ (stock number 8712, Jackson laboratory) B6.FVB-Tg(Acta2- DsRed)1Rkl/J^75^ (stock number 031159, Jackson laboratory), B6.129S-*S*ftpctm1(cre/ERT2)Blh/J^76^ (stock number 028054, Jackson laboratory) were maintained on a C57BL/6N background. For CreER-mediated recombination, mice were injected with Tamoxifen (0.2 mg/g body weight) in peanut oil every other day and these mice were used for experiments three weeks after the last dose. For CCN4 administration, mice were intranasally administered with mouse recombinant CCN4 protein (5 µg/100µl) or PBS 6 times every other day and their lungs were analyzed 3 days after the final dose. For ML-7 injection, mice were intraperitoneally injected with ML-7 (1 mg/ml, 100 µl) or PBS every day after PNX and their lungs were analyzed at 10d. All animal experiments were approved by the Institutional Animal Care and Use Committee of RIKEN Kobe Campus.

### Generation of Fibulin5-/- mouse

Fbln5 knockout mice (*Fbln5*-/-) (accession NO. CDB0249E: https://large.riken.jp/distribution/mutant-list.html) were established with the CRISPR/Cas9-mediated genome editing by zygote electroporation as previously described^77^. The gRNA sequences targeting the 5’ and 3’ region of the mouse *Fbln5* gene were designed to delete exon2∼4 (approximately 6.5 kbp) by using CRISPRdirect^78^ (https://crispr.dbcls.jp/). A new in-frame stop codon emerged in the knockout allele. The insertion of an EcoRI recognition site within the deleted site was mediated by the single strand oligodeoxynucleotides (ssODN). the crRNAs and tracrRNA and the ssODN sequences were as follows; 5’-crRNA (5’- CCA UAU GCU UGC AAG CGA AUG UUU UAG AGC UAU GCU GUU UUG-3’), 3’-crRNA (5’- AGC UAG GCC AUU UCG CAG UUG UUU UAG AGC UAU GCU GUU UUG-3’), tracrRNA (5’- AAA CAG CAU AGC AAG UUA AAA UAA GGC UAG UCC GUU AUC AAC UUG AAA AAG UGG CAC CGA GUC GGU GCU-3’), and ssODN 5’-GTG GGG TTA GAG TAG AGG GCT GGA AGC GCT CTT GGG CTC TGC TGA GTG TGC CCT ATT GAA TTC GTT AGG ATA TGA GAC CAG CCA ACT GCT CCC ATC TAG TAG AAA ATA ATG GTA CCA AAT-3’. The knockout allele was identified by PCR by using following primers; common forward: 5’-CCCAACACATGAAGCAAGTGTGAG-3’, wild type reverse: 5’- GGAAGCCAGAGTGCCAAGATGG-3’ (wild type: 240 bp), and knockout reverse: 5’- GTAACAGGTTCAGAACCACAAGTGG-3’ (knockout: 636 bp).

## Generation of Sox2-P2A-H2B-mCherry reporter mouse

The Sox2-P2A-H2B-mCherry reporter mice (Accession No.CDB0127E: https://large.riken.jp/distribution/mutant-list.html) were generated by CRISPR/Cas9-mediated knockin in zygotes as previously described^79^. For microhomology-mediated end joining (MMEJ)-based knockin, a donor vector containing microhomology arms flanking the P2A-H2B-mCherry cassette^80,81^ was constructed to insert the cassette 3 bp upstream of the PAM site. The crRNA and tracrRNA sequences were as follows: crRNA1 targeting the genomic locus (5′-UGC CCC UGU CGC ACA UGU GAG UUU UAG AGC UAU GCU GUU UUG-3′), crRNA2 targeting the donor vector (5′-GCA UCG UAC GCG UAC GUG UUG UUU UAG AGC UAU GCU GUU UUG-3′), and tracrRNA (5′-AAA CAG CAU AGC AAG UUA AAA UAA GGC UAG UCC GUU AUC AAC UUG AAA AAG UGG CAC CGA GUC GGU GCU-3′). The knockin allele was identified by PCR by using the following primers; forward: 5’- ACCAGCTCGCAGACCTACAT-3’, reverse: 5’-CCCTCCCAATTCCCTTGTAT-3’ (wild type: 387 bp, knockin: 1560 bp).

## Human lung tissues

Human lung tissues were obtained from four patients who received surgery for suspected early-stage lung cancer and did not have apparent emphysematous nor fibrotic changes on their CT images. Only normal lung specimens away from lung cancer lesions were used for analysis. Lung tissues were vigorously washed with sterile PBS and fixed with 4% PFA overnight at 4 °C. Frozen blocks and cryosections were prepared with the same procedures for mouse lungs. The use of human lung samples has been approved by the Institutional Review Board of Kobe University Graduate School of Medicine (B210182). Informed consent has been obtained from all the patients.

## Pneumonectomy (PNX)

PNX was performed as previously described^82^. Briefly mice (3-4 month-old) were anesthetized with isoflurane and intubated using Harvard mini-vent ventilator with 200 µl stroke volume at 200 strokes per minute. The left pulmonary vasculature and main bronchi were ligated with a titanium clip and then the left lobe was resected. The ribs and skin were closed with suture and stainless steel wound clips after removing air to re-establish negative pressure by using a catheter. Buprenorphine (0.1 mg/kg) was injected after surgery. Mice were disconnected from the ventilator when autonomous breathing recovered. Lungs from sham surgery mice were used as controls.

## Immunofluorescence analysis

Lungs were inflated with 4% paraformaldehyde (PFA) to 30 cm H2O pressure and fixed at 4 °C overnight. Lung lobes were separated and washed with PBS for 1 h and then incubated in 30% sucrose/PBS at 4°C for 4 h. Lobes were submerged in 1:1 30% sucrose:Optical Cutting Temperature (OCT) for 8 h followed by incubation in OCT overnight. Lobes were embedded in OCT blocks and cryosectioned at 8-10 µm thickness. Sections were washed with PBS and blocked with 3% BSA, and 0.1% Triton X-100 for 30 min at room temperature. Primary antibodies diluted in the blocking buffer were applied and incubated overnight at 4 °C. Sections were washed with 0.05% Tween20 in PBS and were incubated with fluorophore-conjugated secondary antibodies diluted in the blocking buffer for 1 h at room temperature. DAPI was used to detect nuclei. All primary and secondry antibodies are listed in Reporting Summary. For antibodies recognizing phosphorylated protein (pMLC, pSMAD2/3), PhosStop was added to the fixative. Images were obtained using Zeiss LSM 710 and 910 confocal microscope.

## Proximity ligation in situ hybridization (PLISH)

The PLISH protocol was described previously^83^. Briefly, frozen sections were fixed with 4% PFA for 20 min, treated with 20 µg/ml proteinase K for 8 min at 37 °C, and dehydrated with ethanol. The sections were incubated with gene-specific probes in hybridization buffer (1 M sodium trichloroacetate, 50 mM Tris [pH7.4], 5 mM EDTA, 0.2 mg/ml heparin in diethylpyrocarbonate (DEPC)-treated H2O) for 2 h at 37 °C followed by washing with hybridization buffer. Common bridge and circle probes were added to the sections and incubated for 1 h followed by T4 ligase reaction for 2 h. Rolling circle amplification was performed by using phi29 polymerase overnight. The sections were washed with label probe hybridization buffer (2x saline-sodium citrate/20% formamide in DEPC-H2O) and fluorophore-conjugated detection probes were applied and incubated for 1 h at 37 °C. DAPI was used for nuclear counter staining.

## Whole mount staining

CUBIC-L/RA clearing was used for 2-color staining and endogenous red fluorescence reporter imaging^84^. Tissues were fixed with 4% PFA at least 4 h at 4 °C and washed with PBS, followed by immersion in CUBIC-L (10% N-Butyldiethanolamine, 10% Triton X-100) for 4 d. Tissues were washed with PBS and HEPES-TSC (pH 7.5) (10mM HEPES, 10% Triton X-100, 200 mM NaCl, 0.5% Casein, 0.05% NaN3). A primary antibody and a secondary antibody were mixed in HEPES-TSC for 1.5 h at 37 °C with tapping every 30 min and applied to lung tissues for 4 d at 37 °C with gentle shaking. After washing with 0.1 M PBT and 0.1 M PB, tissues were fixed with 1% formaldehyde overnight at room temperature. Tissues were washed with 0.1 M PB and then immersed in CUBIC-RA (45% Antipyrine, 30% Nicotinamide, 0.5% N-butyldiethanolamine) for 3 d at room temperature for RI matching.

For 3-color staining, BABB (Benzoic Acid Benzyl Benzoate) clearing was used. Sample pretreatment was slightly modified from iDISCO procedures^85^. Tissues were fixed with 4% PFA, washed with PBS, and dehydrated with an up-series of methanol, followed by bleaching with 5% H2O2 in 20% DMSO/methanol at 4 °C overnight. Tissues were washed sequentially with methanol, 20% DMSO in methanol, 80% methanol in PBS, 50% methanol in PBS, PBS, and then PBS containing 0.2% Triton X-100 in order. Tissues were incubated with 0.2 % Triton X-100, 20 % DMSO, 0.3 M glycine in PBS at 37 °C overnight, and then blocked with 0.2 % Triton X-100, 10% DMSO, 6% donkey serum in PBS at 37 °C overnight. Tissues were washed with 0.2 % Tween-20 with 10 µg/ml heparin in PBS (PTwH) and incubated with primary antibodies diluted in PTwH/5 % DMSO/3 % donkey serum at 37 °C for at least 4 d. Tissues were washed with PTwH 5 times and incubated with secondary antibodies diluted in PTwH/3 % donkey serum at 37 °C for 4 d. After washing with PTwH for 1-2 d, tissues were dehydrated with methanol gradient and cleared in BABB (1:2 BA:BB). All cleared samples were imaged with Zeiss 710 confocal microscope, Andor Dragonfly202 spinning-disk confocal microscope or Ultramicroscope Blaze (Miltenyi Biotech).

## 5-ethynyl-2’-deoxyuridine (EdU) incorporation assay

For cell proliferation analysis, mice were intraperitoneally injected with EdU (50 mg/kg) 4 h before fixation. For EdU incorporated cell tracing experiment, pregnant dams were intraperitoneally injected with EdU (50 mg/kg) at E14.5 and fixed embryonic lungs at indicated time points. Click-iT Plus EdU Cell Proliferation Kit for Imaging was used to detect EdU-incorporated cells.

## Quantification of mean linear intercept (MLI)

H&E stained lung sections were imaged by EVIDENT IX83 microscope with X20 objective lens. MLI was automatically measured vertically and horizontally with 200-pixcel intervals by using the ImageJ plugin as previously described^86^. At least 3 representative images were used for quantification from one mouse and biological replicates which were averages of each value from three independent mice at each timepoint were shown on the plots.

## Mouse lung dissociation and FACS

Lung dissociation was described previously^87^. Briefly, lungs were inflated with enzyme solution containing collagenase type I (450 U/ml), dispase (5 U/ml), and DNaseI (0.33 U/ml) in DMEM/F12. Lobes were separated, cut into small pieces and incubated with 3 ml enzyme solution for 30 min at 37 °C with rotation. An equal amount of 10% FBS DMEM/F12 was added to quench the enzymes and filtered through a 100 µm strainer. The cell pellet was resuspended and incubated with 2 ml of red blood cell lysis buffer for 2 min at room temperature. The cell suspension was washed with DMEM/F12, filtered through a 40 µm strainer, and resuspended in 2% BSA-containing DMEM/F12. The cells were stained with antibodies and sorted with BD FACSAria Fusion Flow Cytometer.

## Quantitative Real Time PCR

Cells were lysed with TRIzol Reagent. Total RNA was purified with Direct-zol RNA microprep kit (Zymo Research) according to manufacturer’s instructions. Complimentary DNA library was synthesized from total RNA with SuperScript III First-Strand Synthesis System. Quantitative Real Time PCR was performed by using gene-specific primer sets with Thunderbird SYBR qPCR mix on 7500 Real Time PCR system. PCR cycling parameters were 95 °C for 5 min (one cycle); 95 °C for 15 sec, 62 °C for 15 sec, and 72 °C for 35 sec (40 cycles). *Gapdh* (glyceraldehyde 3-phosphate dehydrogenase) was used for normalization of gene expression as a housekeeping gene. Fold-changes in expression of targeted genes were calculated using the 2-ΔΔCt method. Only biological replicates were shown on the plots and these were an average of more than 3 technical replicates.

## Single cell RNA sequencing

The lungs were dissociated 1 d, 4 d, and 9 d after PNX as described earlier. Mice without surgery were used as a control. Total lung cells from the accessory lobes of two mice were pooled at each time point. The cells were filtered through a 40 µm strainer and then 16,500 cells from each time point were subjected to the library construction by using Chromium Next GEM Single Cell 3’ Reagent Kits v3.1 (Dual Index) according to the manufacturer’s instructions. The average fragment size was determined by Bioanalyzer with Agilent Bioanalyzer High Sensitivity chip for library quality control. Libraries were sequenced on illumina NovaSeq X Plus to obtain a sequencing depth of at least 40,000 reads per cell.

## scRNA-seq analysis

FASTQ files were processed using CellRanger v7.1.0 (10x Genomics) using the mouse mm10 (GENCODE vM23/Ensembl 98, refdata-gex-mm10-2020-A) genome as a reference. Unique molecular identifier barcodes (UMIs) counting and feature-barcode matrix generation were conducted by Cell Ranger count pipeline with default setting, and then background noise such as ambient RNAs and barcode swapping was removed by using CellBender^88^ remove-background function with a value of 5e- 5 for the learning rate. Further downstream analysis such as quality check, normalization, scaling, dimensionality reduction, clustering, and visualization were performed in R using the Seurat^89^ v 4.3.0. Low quality cells which have less than 300 features and more than 10 % mitochondrial count were filtered. Potential doublet cells were removed by using the DoubletFinder package^90^. Total 31,963 cells were obtained through these pre-processing analyses. Principal component analysis (PCA) was performed to reduce dimensionality, and the number of PCs used for downstream analysis was estimated by plotting standard deviations of principal component using the ElbowPlot function. Clustering was performed with a value of 0.5 for resolution and UMAP rendering was used to visualize the clusters. The ductal myofibroblast cluster was identified on the basis of the expression of *Lgr6*, *Hhip*, *Aspn*, and *Cdh4*. For re-analysis of published data, the metadata determined by the original authors were reused. For human gene projection to mouse data, human gene names were converted to corresponding mouse gene names by using org.Hs.eg.db and org.Mm.eg.db as mapping annotation references, and then the module score were calculated by the Seurat AddModuleScore function. The clusterProfiler v4.10.1 was used for GeneOntology and Pathway analysis^91^.

## Gene panel design for Xenium spatial transcriptome analysis

A total of 5,056 genes, including 5,006 genes derived from the pre-designed Xenium Prime 5K Mouse Pan Tissue & Pathway Panel and 50 genes from a custom-designed panel (Panel ID: PX9YD4, list files). The custom genes were selected based on developing mouse lung single-cell RNA-seq data^6^.

## Experimental protocol for Xenium in situ

The lungs were fixed 9d after PNX (n = 2). Formalin-fixed, paraffin-embedded (FFPE) lung blocks were prepared as previously described^92^. Mice normal lung tissue was used as a control (n = 1). FFPE tissue blocks were cut at 5 μm thickness and placed on a Xenium slide (10x Genomics) according to the manufacturer’s protocol (10x Genomics, CG000578). Tissue deparaffinization and decrosslinking steps were conducted according to the manufacturer’s protocol (10x Genomics, CG000580). Subsequently, probe hybridization, ligation, amplification, cell-segmentation staining, autofluorescence quenching and nuclear staining were performed according to the manufacturer’s protocol (10x Genomics, CG000760).

## Xenium data generation

Finalized Xenium slides were loaded onto the Xenium Analyzer instrument, and data were acquired using the onboard software (XOA v3.1.0.4). Using the initial Xenium transcript file, Cell re-segmentation was performed with the transcript-based tool, Proseg (v2.0.2)^93^. Proseg cell-segmentation outputs were subsequently converted into Xenium-compatible outputs using Xenium Ranger (v3.1.1.0).

## Xenium data analysis

Proseg-processed Xenium data were loaded and analyzed with scanpy^94^ (v1.10.3) and squidpy^95^ (v1.6.1). Cells with fewer than 30 counts were excluded. The count data were normalized by total count and log- transformed using the log1p function. The data were subsequently scaled to zero mean and unit variance. A neighborhood graph was calculated with 200 principal components. A batch-balanced neighborhood graph was constructed using BBKNN with sample as the batch key^96^. Three-dimensional UMAP embeddings were generated with default parameters. Leiden clustering^97^ was performed; resolution was 1.5. Clusters were annotated based on with DEGs identified using the Wilcoxon rank-sum test established marker genes. Major cell types were defined as “Endothelial”, “Epithelial”, “Mesenchymal”, “Mesothelial”, “Immune”, “Proliferative”.

Cells classified as “Proliferative” cluster was re-clustered as above and reassigned to cell type categories. Following this reassignment, the three major populations, “Endothelial”, “Epithelial”, “Mesenchymal”, were further clustered without batch correction. Clusters were annotated with DEGs and established marker genes. Clusters with low average counts or highly expressing marker genes from other major cell types, suggestive of contamination, were excluded from downstream analyses. During mesenchymal sub-clustering, the cluster characterized by high *Hhip* and *Aspn* expression was annotated as ductal myofibroblasts.

For Xenium analysis, “*Hhip+* Mesenchymal cell-specific genes were calculated with Wilcoxon rank-sum test among “Mesenchymal”. The genes were filtered with log2 fold change >2 and a p-value < 1 × 10⁻⁵. The spatial distribution of annotated ductal myofibroblast in PNX sections was visualized using Xenium Explorer (v3.2.0)

## Embryonic lung explant culture

The lungs dissected from E11.5 embryos were transferred onto Millicell cell culture insert hydrophilic polytetrafluoroethylene membrane (Merck, PICM0RG50) and cultured at an air-liquid interface with DMEM/F12 containing 10% FBS, GlutaMAX, and penicillin/streptomycin. To assess the role of TGF-β signaling in SMA+ cell differentiation, 10 µM of TGF-β inhibitor SB431542 was added to the medium. The explants were analyzed 2 d after culturing.

## RNAscope

RNA in situ hybridization was performed on 10 µm frozen sections using the RNAScope Multiplex Fluorescent Assay v according to the manufacturer’s instructions (Advanced Cell Diagnostics). Tissue sections were dried up at room temperature and washed with H2O, followed by target retrieval for 5 min at 98 °C. After wash and dehydration, tissue sections were incubated with Protease III for 15 min at 40 °C. Slides were hybridized with target probes(Hs-ACTA2-O1 (444771), Hs-LGR6-C2 (410461-C2)) for 2 h at 40 °C. Following wash and signal amplification steps were conducted according to the manufacturer’s instructions.

## QUANTIFICATION AND STATISTICAL ANALYSIS

All results were presented as mean ± standard error from at least three independent experiments. Statistical analyses were performed with R package tidyplot using unpaired Student’s t test (two-tailed) for comparison between two groups or using one-way analysis of variance with Tukey’s Honestly Significant Difference post hoc test for comparisons between more than two groups. P values were provided in the Source Data file and differences with P<0.05 were considered significant. For cell count quantification, at least more than 5 images from one mouse were used for count and all data from at least three independent mice were used for statistics calculations. The representative images were shown in each figure.

## Data availability

The authors declare that all data supporting the findings of this study are available within the article and its supplementary materials, including Source Data. The scRNA-seq datasets and Xenium datasets will be uploaded to GEO database and will be available when the manuscript is accepted. The raw images will be available in the System Science of Biological Dynamics (SSBD) repository when the manuscript is accepted.

## Acknowledgements

We thank the members of Morimoto lab and Konagaya lab for fruitful discussion, Hironobu Fujiwara and Shimpei Gotoh for critical comments on the manuscript. We also thank Brigid Hogan and Purushothama Rao Tata for kindly providing *Pdgfra-CreERT2* mice, Pierre Chambon for kindly providing *SMA-CreER*, Robert Mecham for kindly providing anti-Elastin antibody, Hironobu Fujiwara for kindly providing a mouse line, Atsuyasu Sato and Itsuki Yuasa for technical support on PNX surgery, Laboratory for Animal Resources and Genetic Engineering for animal experiments, Riken Kobe Bioimaging Facilities & Factory for microscope experiments, Genomics Research and Analysis Support Team at RIKEN-BDR for scRNA- seq library preparation and computational analysis. Some schematic diagrams were created with BioRender.com. Some graphs were created with tidyplots^98^.

## Author contributions

H.K. and M.M. conceived the project. H.K. and A.Y. performed experiments and analyzed data. N.N., J.K., D.C., M.K., and T.K. performed and analyzed Xenium spatial transcriptomic experiments. O.N. supported scRNA-seq analysis. H.K., T.A. and H. Kiyonari generated Fbln5 KO and Sox2-P2A-H2B- mcherry mice. D.H., S.T., Y.M., and T.N. obtained human lung specimens. H.K. and M.M. wrote the manuscript. All authors reviewed the manuscript and approved the submission.

## Funding

This research was supported by the Japan Society for the Promotion of Science Grant-in-Aid for Scientific Research 21K16127 (H.K), 25K11444 (H.K), 22H05640(M.M), 24H01415 (M.M), AMED-CREST under 24028395 (H.K, M.M), the RIKEN BDR-Otsuka Pharmaceutical Collaboration Center supporting grant Kakehashi (H.K), and RIKEN internal grant.

## Declaration of interests

The authors declare no competing interests.

## Supplemental information

### Supplemental Fig

**Supplementary Fig. 1.**
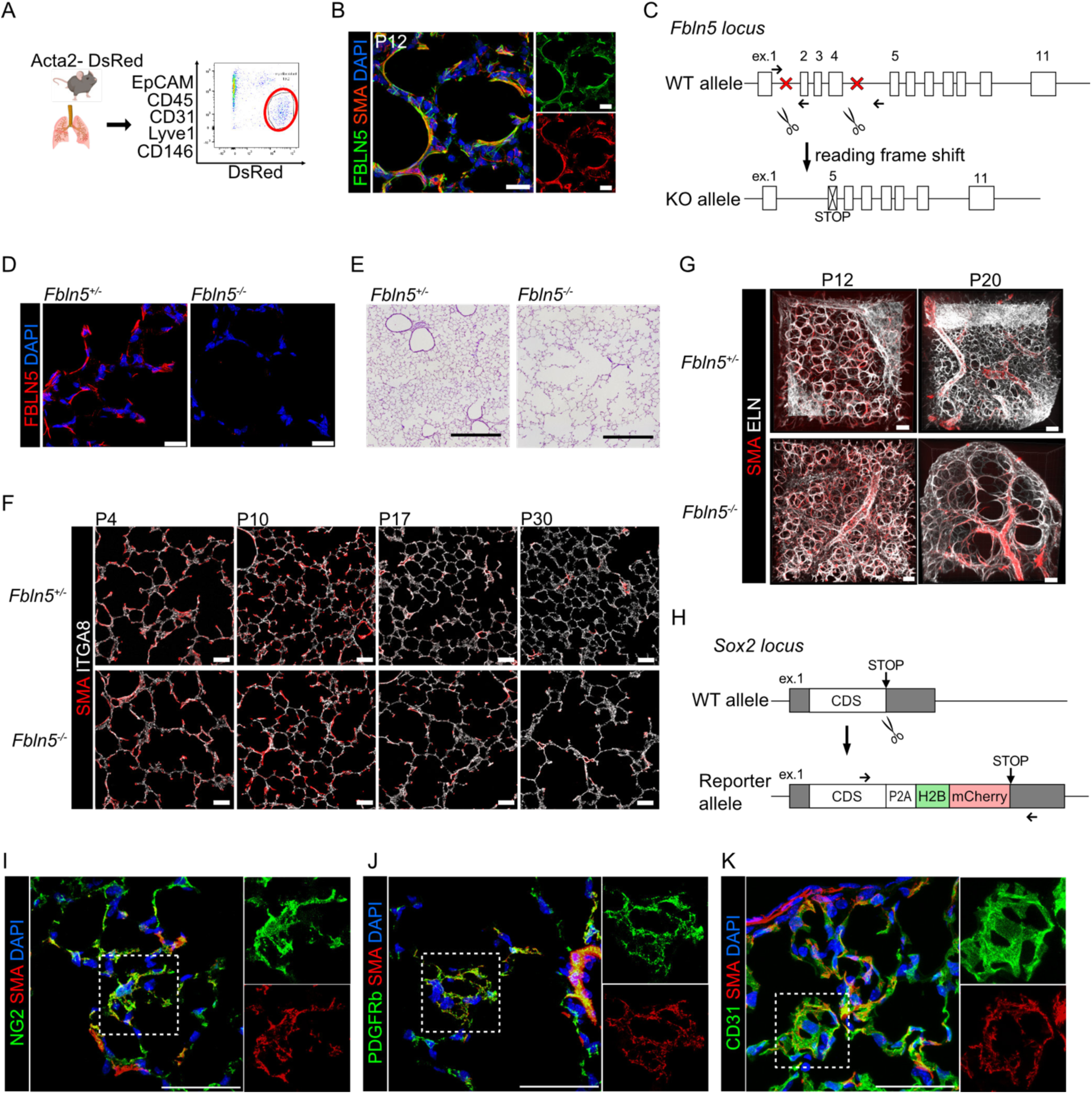
Analysis of postnatal lungs of Fbln5-deficient mice. A. FACS gating strategy for alveolar myofibroblasts from Acta2-DsRed reporter mice. The red circle indicates alveolar myofibroblast population. B. Immunofluorescence images of section stained for FBLN5 (green) and SMA (red) at P12. Scale bars, 20µm. C. Targeting strategy for Fbln5 KO mice. Arrows indicate genotyping primers. D. Immunofluorescence images of lung sections stained for FBLN5 (red) in Fbln5+/- (left) and Fbln5 -/- (right) mice. Scale bars, 20 µm. E. Representative images of H&E staining of Fbln5+/- (left) and Fbln5-/- (right) lungs in adulthood. Scale bars, 500 µm. F. Immunofluorescence images of lung sections stained for SMA (red) and ITGA8 (white) from Fbln5+/- (top) and Fbln5 -/- (bottom) mice. Scale bars, 50 µm. G. Three-dimensional images of cleared lung tissue from *Fbln5*+/- (top) and *Fbln5*-/- (bottom) mice stained for SMA (red) and ELN (white). Scale bars, 30 µm. H. Targeting strategy for Sox2-H2B-mCherry reporter mice. Arrows indicate genotyping primers. I. Immunofluorescence images of lung section stained for NG2 (green) and SMA (red) at PNX 10d. High- magnification images of the insets drawn by white squares (right). Scale bars, 50 µm. J.Immunofluorescence images of lung section stained for PDGFRb (green) and SMA (red) at PNX 10d. High- magnification images of the insets drawn by white squares (right). Scale bars, 50 µm. K. Immunofluorescence images of lung section stained for CD31 (green) and SMA (red) at PNX 10d. High- magnification images of the insets drawn by white squares (right). Scale bars, 50 µm.

**Supplementary Fig. 2.**
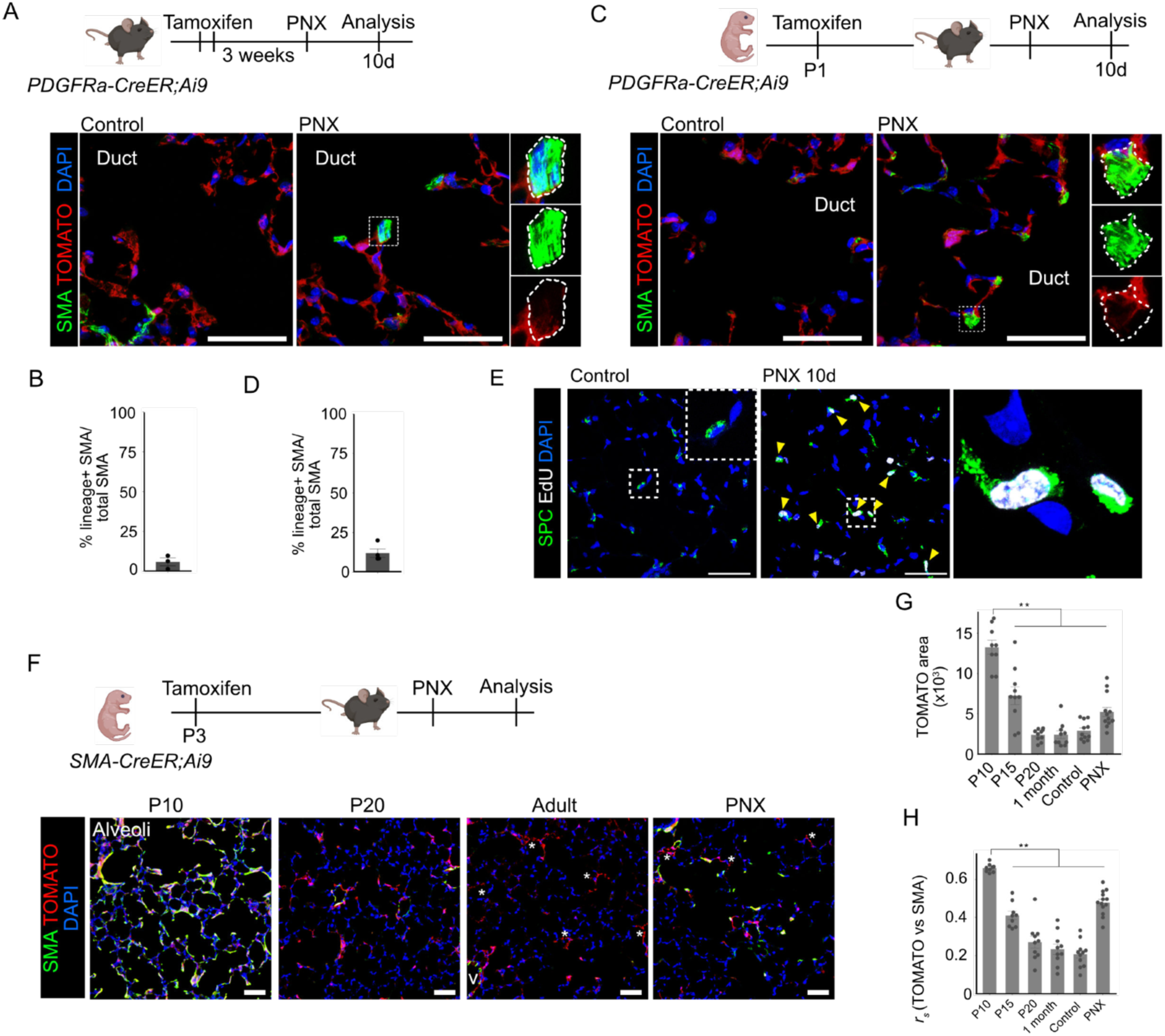
Cellular responses during lung regrowth after PNX. A.Schematic diagram of lineage-tracing experiment of *Pdgfra*+ fibroblasts in adult lungs and immunofluorescence images of sections of the alveolar ducts from control (left) and PNX 10d (right) stained for SMA (green) with signals from TOMATO (red). High-magnification images of the insets drawn by white squares. Three SMA+ cells are outlined with white dashed lines. Scale bars, 50 µm. B. Quantification of the ratio of the *Pdgfra*-lineage+ SMA+ cells in total SMA+ cells in the alveolar ducts after PNX. (n = 3 mice/group) C. Schematic diagram of lineage-tracing experiment of *Pdgfra*+ cells from neonate and immunofluorescence images of sections of the alveolar ducts from control (left) and PNX 10d (right) stained for SMA (green) with signal from TOMATO (red). High-magnification images of the insets drawn by white squares. Three SMA+ cells are outlined with white dashed lines. Scale bars, 50 µm. D. Quantification of the ratio of the Pdgfra-lineage+ cells labeled from neonate in total SMA+ cells in the alveolar ducts after PNX. (n = 4 mice/group) E. Immunofluorescence images of lung sections from control (left) and PNX 10d (right) stained for SPC (green) and EdU (white). High-magnification images of the insets drawn by white squares and arrowheads indicate EdU+ SFTPC+ proliferating AT2 cells. Scale bars, 50 µm. F. Schematic diagram of lineage-tracing experiment of SMA+ cells from neonate and immunofluorescence images of alveolar regions at indicated time points stained for SMA (green) with signal from TOMATO (red). High-magnification images of the insets drawn by white squares. Asterisks indicate pericytes. Scale bars, 50 µm. G. Quantification of the TOMATO+ area in alveoli. Each dot indicates the values from each image. H. Bar graph indicating the co-localization (rs, correlation coefficient) of TOMATO and SMA signals. Each dot indicates the values from each image. (n = 3 mice/group)

**Supplementary Fig. 3.**
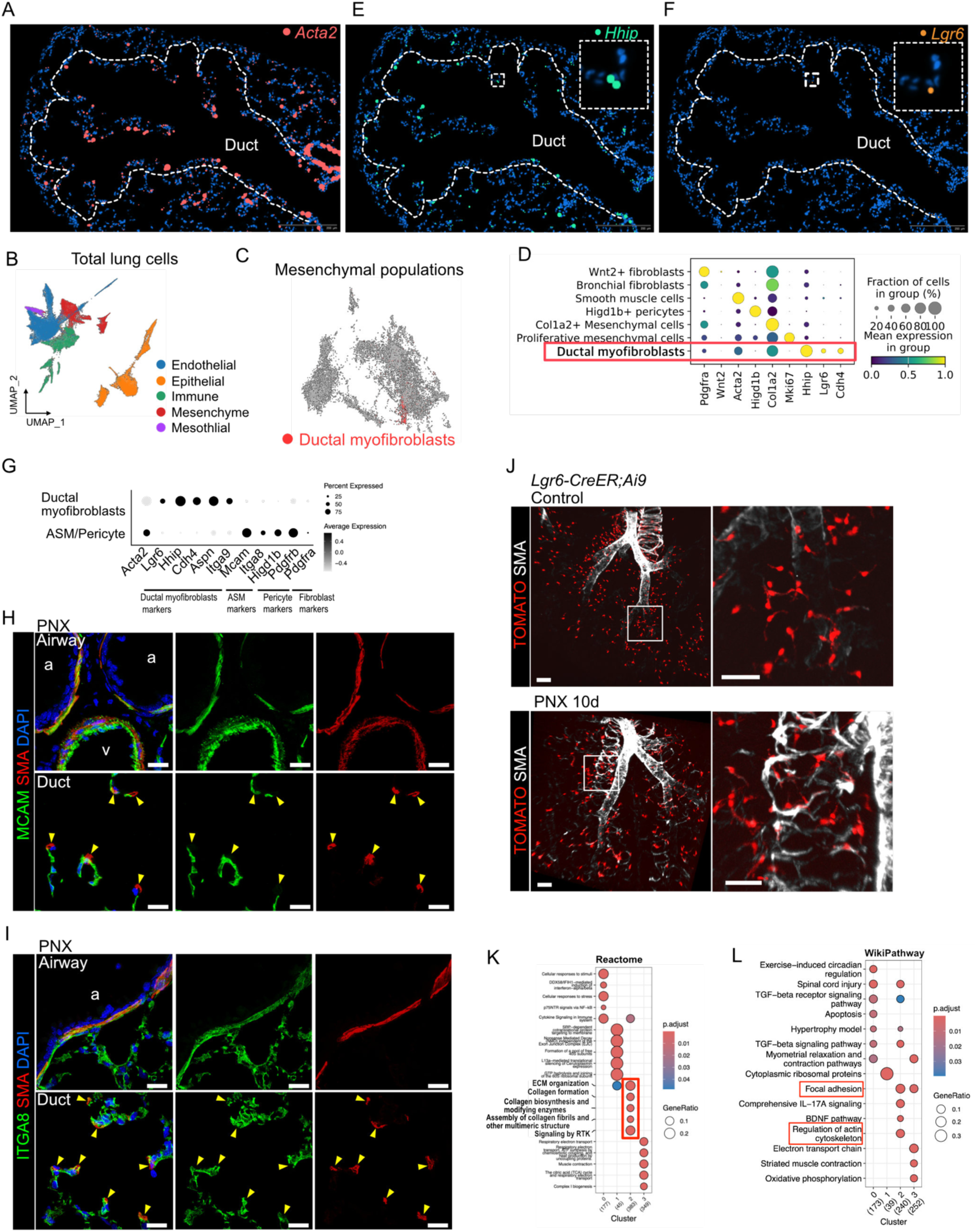
Spatial transcriptomic analysis identified ductal myofibroblasts in adult lungs after PNX. A. Spatial profiling of *Acta2* signals in PNX lung. B. UMAP visualization of major lung cell types by Xenium spatial analysis. C. UMAP plot showing ductal myofibroblast cluster highlighted in red in mesenchymal populations. D. Dot plot showing cluster-specific marker genes in mesenchymal subclusters. Red rectangle shows ductal myofibroblast population. E. Spatial profiling of *Hhip* signals in PNX lung. F. Spatial profiling of *Lgr6* signals in PNX lung. White dashed line indicates alveolar duct. G. Dot plot showing the ductal myofibroblast- and ASM/pericyte-specific markers. H. Immunofluorescence images of the airway (top) and the alveolar duct (bottom) stained for MCAM (green) and SMA (red). Arrowheads indicate MCAM- SMA+ ductal myofibroblasts. Scale bars, 20 µm. I. Immunofluorescence images of the airway (top) and the alveolar duct (bottom) stained for ITGA8 (green) and SMA (red). Arrowheads indicate ITGA8- SMA+ ductal myofibroblasts. Scale bars, 20 µm. J. Three-dimensional images of cleared lung tissues from control (top) and PNX 10d (bottom) of *Lgr6*- CreER;Ai9 mice stained for SMA (white) with signal from TOMATO endogenous fluorescence reporter (red). High-magnification images of the insets drawn by white squares (right). Scale bars, 100 µm (left), 50 µm (right). K. Reactome pathways enriched in each cluster. The red square indicates ECM-related terms. L. WikiPathways enriched in each cluster. The red squares indicate terms related to contractile forces.

**Supplementary Fig. 4.**
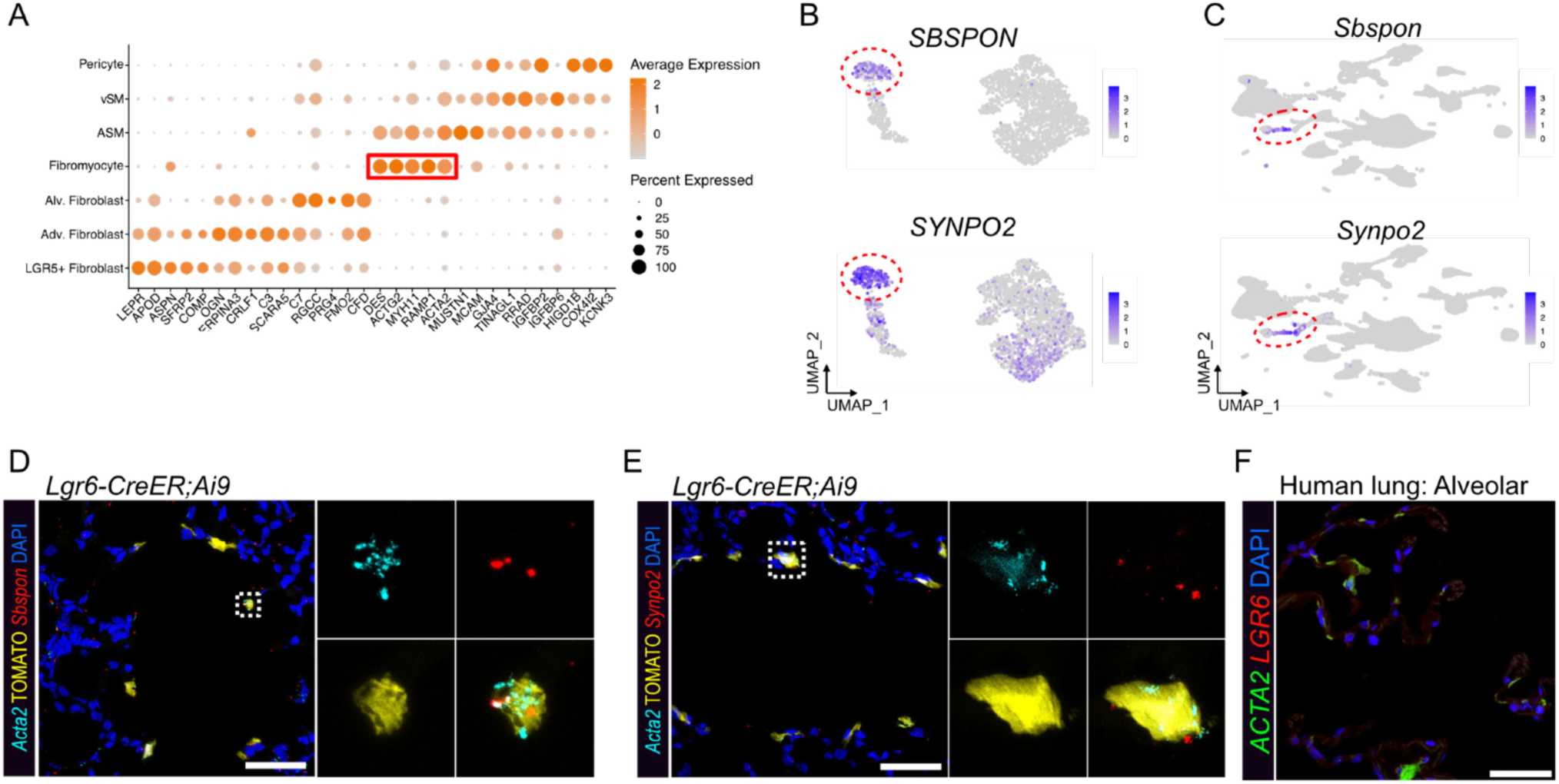
Human fibromyocytes share specific marker gene expression with mouse ductal myofibroblasts. A. Dot plot showing cluster-specific marker gene expression in distal airway mesenchyme data. The annotation was corresponding to Fig. 4A. The red rectangle indicates fibromyocyte-specific genes. B. Feature plots showing the expression of indicated genes in human distal mesenchyme data. C. Feature plots showing the expression of indicated genes in mouse total lung cells. D. Detection of transcripts of *Acta2* (cyan) and *Sbspon* (red) with signal from TOMATO in lungs from *Lgr6*- CreER;Ai9 mice 10 days after PNX. High-magnification images of the insets drawn by white squares (right). Scale bar, 50 µm. E. Detection of transcripts of *Acta2* (cyan) and *Synpo2* (red) with signal from TOMATO in lungs from *Lgr6*- CreER;Ai9 mice 10 days after PNX. High-magnification images of the insets drawn by white squares (right). Scale bar, 50 µm. F. Detection of transcripts of *ACTA2* (green) and *LGR6* (red) in the alveoli in human lungs. Scale bar, 50µm.

**Supplementary Fig. 5.**
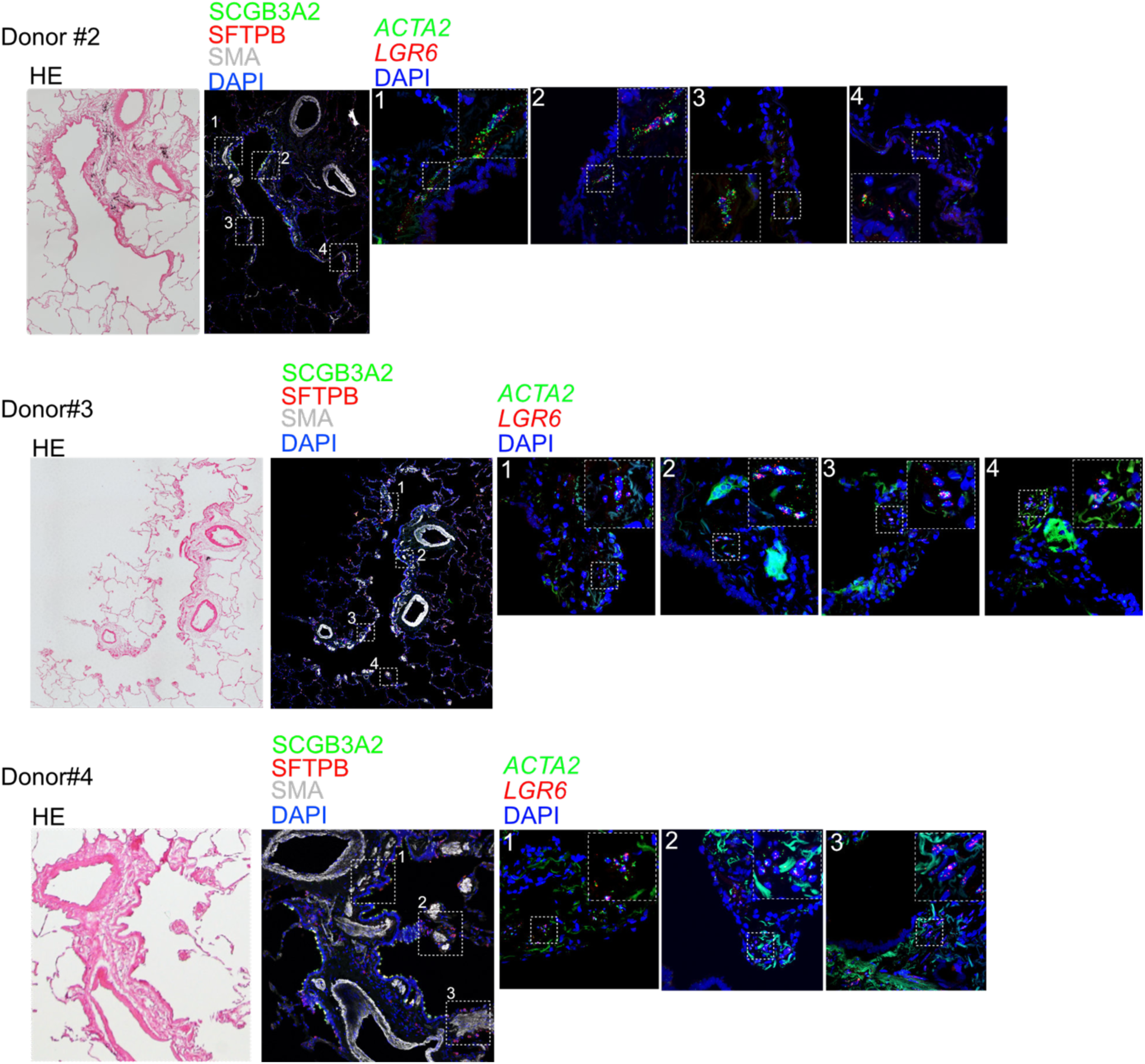
Identification of LGR6+ fibromyocytes in human lungs. Representative images of H&E staining of TRB region from three different donors and immunofluorescence analysis for SCGB3A2 (green), SFTPB (red), and SMA (white) in the same area. In situ hybridization of ACTA2 (green) and LGR6 (red) identified fibromyocytes.

**Supplementary Fig. 6.**
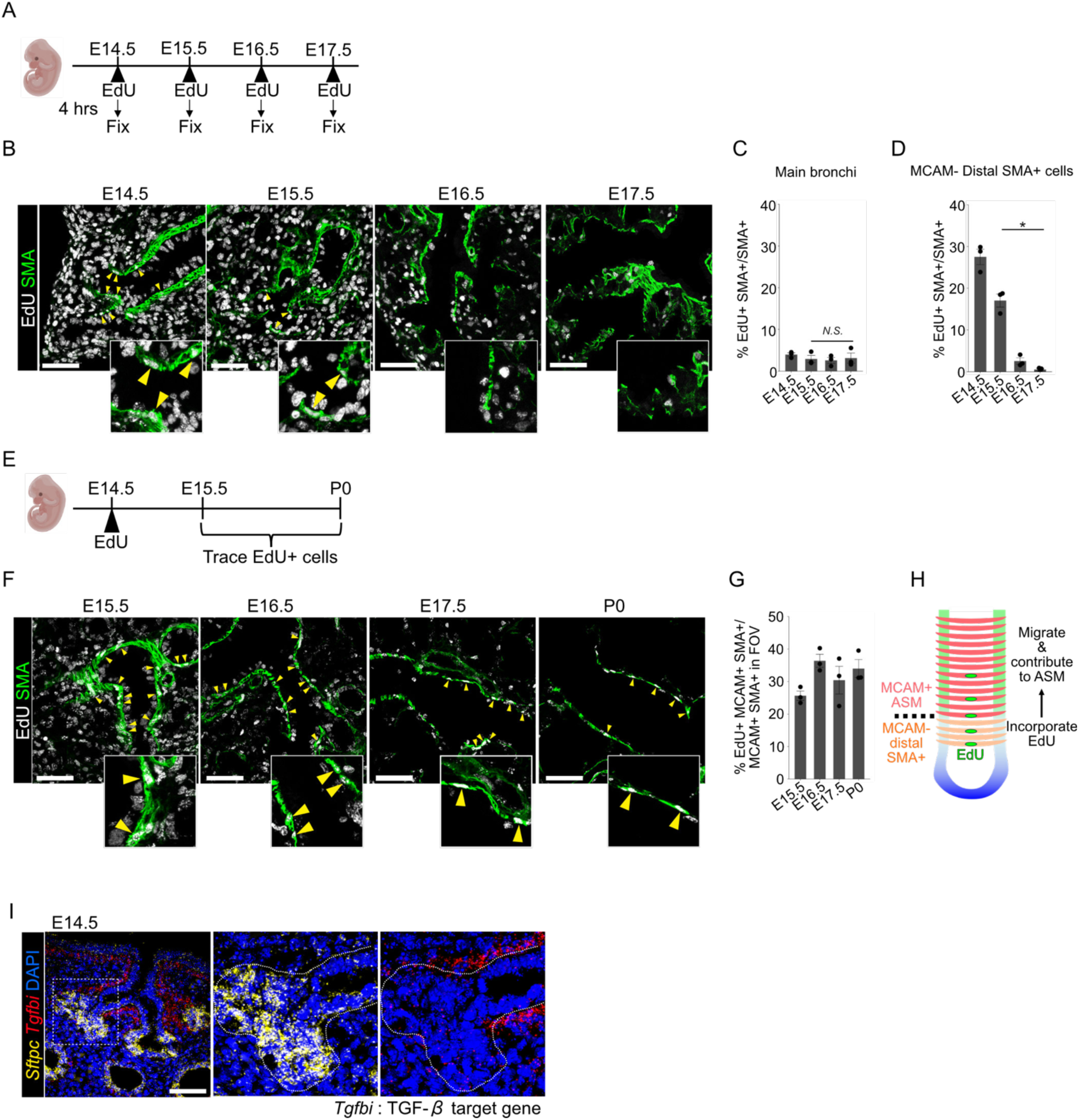
MCAM- SMA+ cells at the distal end function as immediate progenitors of distal airway smooth muscle cells. A. Schematic diagram of EdU incorporation assay in embryo. B. Immunofluorescence images of lung sections stained for SMA (green) and EdU (white) with high-magnification images of the distal end of SMA+ bundles. Yellow arrowheads indicate EdU+ proliferating SMA+ cells. Scale bars, 50 µm. C. Quantification of the ratio of EdU+ SMA+ cells in total SMA+ cells in main bronchi. D. Quantification of the ratio of EdU+ SMA+ cells in total SMA+ cells in MCAM- distal SMA+ population. E. Schematic diagram of EdU-tracing experiment. F. Immunofluorescence images of lung sections stained for SMA (green) and EdU (white) with high magnification images of EdU+ SMA+ cells (yellow arrowheads). Scale bars, 50 µm. G. Quantification of the ratio of EdU+ MCAM+ SMA+ cells in total MCAM+ SMA+ airway smooth muscle population. H. Summary of the EdU-incorporation and tracing experiments. I. Detection of transcripts of *Sftpc* (yellow) and *Tgfbi* (red) in E14.5 lungs. Scale bars, 50 µm. Data represents the mean ± SEM obtained from three independent mice.

**Table S1.**
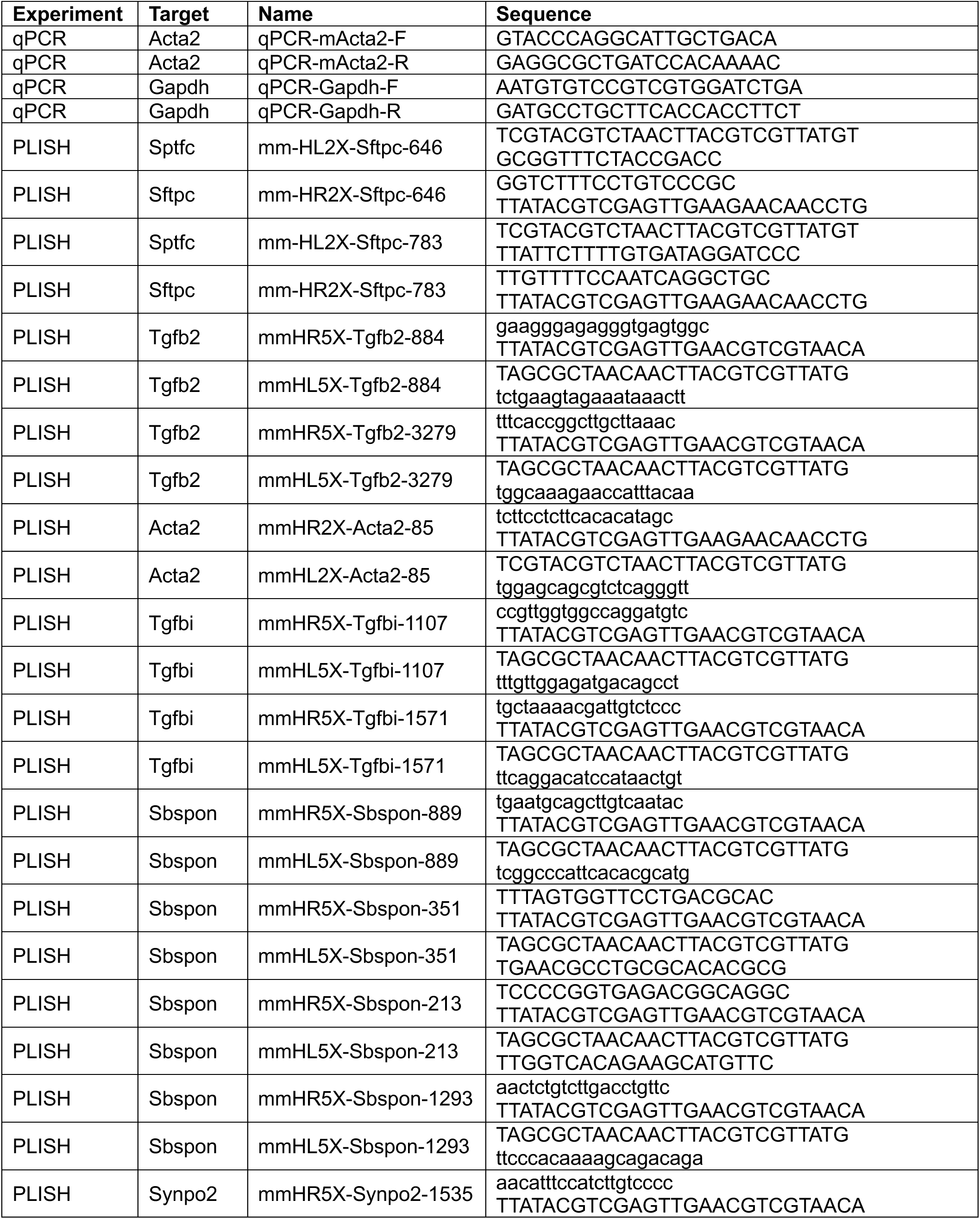
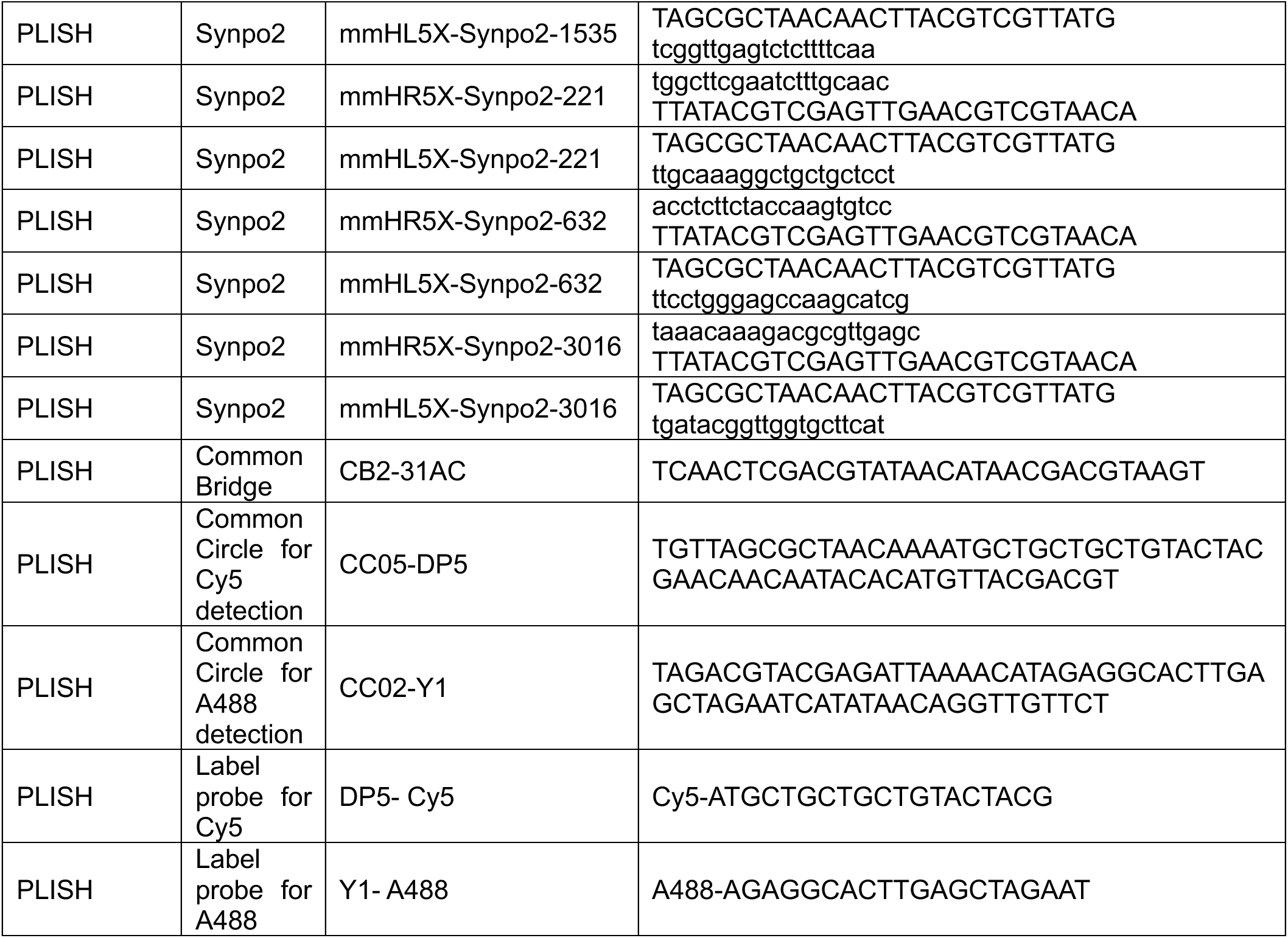

## Notes

### Competing Interest Statement

The authors have declared no competing interest.

